## Supplementary Materials for "Neural Correlates and Reinstatement of Recent and Remote Memory: A Comparison Between Children and Young Adults"

### S1. Supplementary Methods

#### S1.1. Assessment of demographic and cognitive covariates

Other cognitive covariate tasks, such as cognitive switching and object-location memory, were run on each session but they are not included in the current paper.

***Day 0*:** After the experimental task, several subtests of the K-ABC II Test *(e.g., Atlantis, Rover, Rebus, Riddle and Atlantis delayed) were administered to children, while young adults were tested with the WAIS-IV Test.

***Day 1:*** In addition, children performed several subtests of the K-ABC II Test *(e.g., Expressive Vocabulary, Triangles, Pattern Reasoning), and a cognitive switching task.

**Day 14:** Children performed several subtests of the K-ABC II Test *(e.g., Patterns, Verbal Knowledge, Word Order), and an object-location memory task.

In addition to the experimental paradigm, a sleep diary to assess the quality and duration of sleep was completed daily for the 14-day period between learning and long-delay.

#### S1.2. FMRI data pre-processing

**The following description of the fMRI data pre-processing was generated by fMRIPrep 22.0.0:**

Results included in this manuscript come from preprocessing performed using fMRIPrep 22.0.0 (Esteban et al., 2018, 2019; RRID:SCR_016216), which is based on Nipype 1.8.3 (Gorgolewski et al., 2011; Gorgolewski et al., 2016); RRID:SCR_002502).

##### *S1.2.1.Preprocessing of B_0_* *inhomogeneity mappings*

A total of 2 fieldmaps were found available within the input BIDS structure for this particular subject. A B_0_-nonuniformity map (or fieldmap) was estimated based on two (or more) echo-planar imaging (EPI) references with topup (Andersson et al. (2003); FSL 6.0.5.1:57b01774).

##### *S1.2.2. Anatomical data preprocessing*

A total of 2 T1-weighted (T1w) images were found within the input BIDS dataset. All of them were corrected for intensity non-uniformity (INU) with N4BiasFieldCorrection (Tustison et al., 2010), distributed with ANTs 2.3.3 (Avants et al. (2008); RRID:SCR_004757). The T1w-reference was then skull-stripped with a Nipype implementation of the antsBrainExtraction.sh workflow (from ANTs), using OASIS30ANTs as target template. Brain tissue segmentation of cerebrospinal fluid (CSF), white-matter (WM) and gray-matter (GM) was performed on the brain-extracted T1w using fast (FSL 6.0.5.1:57b01774, RRID:SCR_002823; Zhang et al., (2001)). A T1w-reference map was computed after registration of 2 T1w images (after INU-correction) using mri_robust_template (FreeSurfer 7.2.0; Reuter et al., (2010)). Volume-based spatial normalization to two standard spaces (MNI152NLin6Asym, MNI152NLin2009cAsym) was performed through nonlinear registration with antsRegistration (ANTs 2.3.3), using brain-extracted versions of both T1w reference and the T1w template. The following templates were selected for spatial normalization: FSL’s MNI ICBM 152 non-linear 6th Generation Asymmetric Average Brain Stereotaxic Registration Model [Evans et al. (2012); RRID:SCR_002823; TemplateFlow ID: MNI152NLin6Asym], ICBM 152 Nonlinear Asymmetrical template version 2009c [ Fonov et al. (2009); RRID:SCR_008796; TemplateFlow ID: MNI152NLin2009cAsym].

##### *S1.2.3. Functional data preprocessing*

For each of the 5 BOLD runs found per subject (across all tasks and sessions), the following preprocessing was performed. First, a reference volume and its skull-stripped version were generated by aligning and averaging 1 single-band references (SBRefs). Head-motion parameters with respect to the BOLD reference (transformation matrices, and six corresponding rotation and translation parameters) are estimated before any spatiotemporal filtering using mcflirt (FSL 6.0.5.1:57b01774; Jenkinson et al. (2002)). The estimated fieldmap was then aligned with rigid-registration to the target EPI (echo-planar imaging) reference run. The field coefficients were mapped on to the reference EPI using the transform. BOLD runs were slice-time corrected to 0.346s (0.5 of slice acquisition range 0s-0.693s) using 3dTshift from AFNI ( Cox & Hyde, (1997); RRID:SCR_005927). The BOLD reference was then co-registered to the T1w reference using mri_coreg (FreeSurfer) followed by flirt (FSL 6.0.5.1:57b01774; Jenkinson & Smith (2001) with the boundary-based registration (Greve & Fischl, 2009) cost-function. Co-registration was configured with six degrees of freedom. First, a reference volume and its skull-stripped version were generated using a custom methodology of fMRIPrep. Several confounding time-series were calculated based on the preprocessed BOLD: framewise displacement (FD), DVARS and three region-wise global signals. FD was computed using two formulations following Power (absolute sum of relative motions, Power et al. (2014) and Jenkinson et al. (2002) (relative root mean square displacement between affines). FD and DVARS are calculated for each functional run, both using their implementations in Nipype (following the definitions by Power et al. (2014)). The three global signals are extracted within the CSF, the WM, and the whole-brain masks. Additionally, a set of physiological regressors were extracted to allow for component-based noise correction (CompCor; Behzadi et al. (2007)). Principal components are estimated after high-pass filtering the preprocessed BOLD time-series (using a discrete cosine filter with 128s cut-off) for the two CompCor variants: temporal (tCompCor) and anatomical (aCompCor). tCompCor components are then calculated from the top 2% variable voxels within the brain mask. For aCompCor, three probabilistic masks (CSF, WM and combined CSF+WM) are generated in anatomical space. The implementation differs from that of Behzadi et al. in that instead of eroding the masks by 2 pixels on BOLD space, a mask of pixels that likely contain a volume fraction of GM is subtracted from the aCompCor masks. This mask is obtained by thresholding the corresponding partial volume map at 0.05, and it ensures components are not extracted from voxels containing a minimal fraction of GM. Finally, these masks are resampled into BOLD space and binarized by thresholding at 0.99 (as in the original implementation). Components are also calculated separately within the WM and CSF masks. For each CompCor decomposition, the k components with the largest singular values are retained, such that the retained components’ time series are sufficient to explain 50 percent of variance across the nuisance mask (CSF, WM, combined, or temporal). The remaining components are dropped from consideration. The head-motion estimates calculated in the correction step were also placed within the corresponding confounds file. The confound time series derived from head motion estimates and global signals were expanded with the inclusion of temporal derivatives and quadratic terms for each (Satterthwaite et al., 2013). Frames that exceeded a threshold of 0.5 mm FD or 1.5 standardized DVARS were annotated as motion outliers. Additional nuisance timeseries are calculated by means of principal components analysis of the signal found within a thin band (crown) of voxels around the edge of the brain, as proposed by Patriat et al. (2017). The BOLD time-series were resampled into several standard spaces, correspondingly generating the following spatially-normalized, preprocessed BOLD runs: MNI152NLin6Asym, MNI152NLin2009cAsym. First, a reference volume and its skull-stripped version were generated using a custom methodology of fMRIPrep. Automatic removal of motion artifacts using independent component analysis (ICA-AROMA; Pruim et al. (2015)) was performed on the preprocessed BOLD on MNI space time-series after removal of non-steady state volumes and spatial smoothing with an isotropic, Gaussian kernel of 6mm FWHM (full-width half-maximum). Corresponding “non-aggresively” denoised runs were produced after such smoothing. Additionally, the “aggressive” noise-regressors were collected and placed in the corresponding confounds file. All resamplings can be performed with a single interpolation step by composing all the pertinent transformations (i.e. head-motion transform matrices, susceptibility distortion correction when available, and co-registrations to anatomical and output spaces). Gridded (volumetric) resamplings were performed using antsApplyTransforms (ANTs), configured with Lanczos interpolation to minimize the smoothing effects of other kernels (Lanczos, 1964). Non-gridded (surface) resamplings were performed using mri_vol2surf (FreeSurfer). Many internal operations of fMRIPrep use Nilearn 0.9.1 ( Abraham et al. (2014); RRID:SCR_001362), mostly within the functional processing workflow.

### S2. Supplementary behavioural results

Table S1

*Statistical overview of the linear mixed effects model for memory retention rates for initially correctly learned items (corrected for chance performance).*

|  | (A) **Recent Memory Retention** | | | **(B) Overall Memory Retention** | |
| --- | --- | --- | --- | --- | --- |
| *Predictors* | *F-value_(DenDF)_* | | *p-value* | *F-value_(DenDF)_* | *p-value* |
|  | 5.19_(1,75)_  47.44_(1,83)_  2.39_(1,88)_  1.73_(1,89)_  1.77_(1,75)_ | |  |  |  |
| Session |  |  | **.026** |  |  |
| Group |  |  | **<.001** | 55.00_(1,85)_ |  |
| Item Type |  |  |  | 229.17_(3,250)_ | **<.001** |
| IQ |  |  | .125 | 5.82_(1,86)_ | **.018** |
| Sex |  |  | .191 | 2.57_(1,87)_ | .113 |
| Session x Group |  |  | .187 |  |  |
| Item Type x Group |  | |  | 17.35_(3,250)_ | **<.001** |
| **Random Effects** |  |  |  |  |  |
| σ^2^ | 59.91 |  |  | 57.36 |  |
| τ_00_ _subNo_ | 74.73 |  |  | 26.37 |  |
| ICC | .56 |  |  | .31 |  |
| N _subNo_ | 88 |  |  | 88 |  |
| Observations | 158 |  |  | 336 |  |
| Marginal R^2^ / Conditional R^2^ | 0.335/ 0.704 |  |  | .659/.767 |  |

*Notes.* Subject was included as random intercept. Group (children and young adults), Session (Day 1, Day 14 and Day1 and Day14 _after 30 minutes_), Item Type (baseline_learning_, immediate, recent vs remote) were included as fixed effects. IQ, Sex, Handedness were included as covariates. ^a^The following reference levels where used: for Session, Day 1/14; for Group, Children; for Item Type, baseline; for Sex, male; for Handedness, right-side handedness. IQ = Intelligence Quotient; σ2 – residuals, τ00 – variance of the random intercept. Type III Analysis of Variance Table with Satterthwaite's method.

Table S1.1

*Statistical overview of the linear mixed effects model for memory retention rates for initially correctly learned items (corrected for chance performance) based on participants who needed only two learning cycles (N = 28).*

|  | (A) **Recent Memory Retention** | | | **(B) Overall Memory Retention** | |
| --- | --- | --- | --- | --- | --- |
| *Predictors* | *F-value_(DenDF)_* | | *p-value* | *F-value_(DenDF)_* | *p-value* |
|  | 4.64_(1,55)_  30.16_(1,59)_  .02_(1,60)_  4.01_(1,60)_  1.18_(1,55)_ | |  |  |  |
| Session |  |  | **.035** |  |  |
| Group |  |  | **<.001** | 33.53_(1,65)_ | **<.001** |
| Item Type |  |  |  | **230.02** _(3,192)_ | **<.001** |
| IQ |  |  | .885 | 1.49_(1,66)_ | **.226** |
| Sex |  |  | .049 | 4.42_(1,66)_ | .**039** |
| Session x Group |  |  | .281 |  |  |
| Item Type x Group |  | |  | 10.56_(3,192)_ | **<.001** |
| **Random Effects** |  |  |  |  |  |
| σ^2^ | 32.92 |  |  | 42.11 |  |
| τ_00_ _subNo_ | 29.74 |  |  | 16.45 |  |
| ICC | .47 |  |  | .28 |  |
| N _subNo_ | 67 |  |  | 67 |  |
| Observations | 122 |  |  | 258 |  |
| Marginal R^2^ / Conditional R^2^ | 0.331/ 0.649 |  |  | .697/.782 |  |

*Notes.* Subject was included as random intercept. Group (children and young adults), Session (Day 1, Day 14 and Day1 and Day14 _after 30 minutes_), Item Type (baseline_learning_, immediate, recent vs remote) were included as fixed effects. IQ, Sex, Handedness were included as covariates. ^a^The following reference levels where used: for Session, Day 1/14; for Group, Children; for Item Type, baseline; for Sex, male; for Handedness, right-side handedness. IQ = Intelligence Quotient; σ2 – residuals, τ00 – variance of the random intercept. Type III Analysis of Variance Table with Satterthwaite's method.

Table S1.2

*Statistical overview of post hoc analysis of the Item Type x Group Interaction effects for the linear mixed effects model for memory retention rates for initially correctly learned items (corrected for chance performance) based on participants who needed only two learning cycles.*

| Contrast | Estimate | df | t | p-value |
| --- | --- | --- | --- | --- |
| d0 vs d1/14 recent CH vs. YA | 10.86 | 197 | 4.701 | <.001 |
| d0 vs d1 remote CH vs. YA | 8.67 | 201 | 3.720 | .003 |
| d0 vs d14 remote CH vs. YA | 11.41 | 201 | 4.736 | <.001 |
| d1/14 vs d1/14 CH | 14.857 | 216.182 | 8.820 | .000 |
| d1/14 vs d1/14 YA | 1.913 | 216.182 | 1.199 | .968 |
| d0 vs d1/d14 recent CH | 12.77 | 197 | 7.248 | <.001 |
| d0 vs d1/d14 recent YA | 1.91 | 197 | 1.28 | .946 |
| d0 vs d1 remote CH | 14.00 | 198 | 7.859 | <.001 |
| d0 vs d1 remote YA | 5.33 | 198 | 3.53 | .006 |
| d0 vs d14 remote CH | 35.75 | 201 | 19.34 | <.001 |
| d0 vs d14 remote YA | 24.4 | 200 | 15.77 | <.001 |

*Notes.* d – Day; CH – children; YA – young adults; df – degrees of freedom, t – t-test; All post hoc test were Sidak corrected.

Figure S1

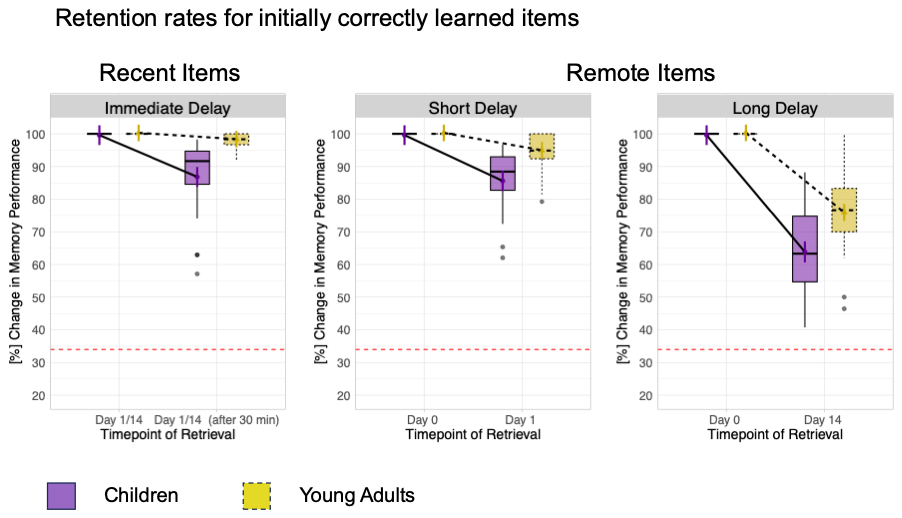

**Retention rates for initially correctly learned items (for participants who needed only two learning cycles).** Memory accuracy is operationalized as the percentage of correct responses in the retrieval task conducted during the MRI scanning sessions for items that were initially correctly learned, indicating initially strong memories. Memory accuracy for recently consolidated items did not differ between sessions in young adults and children and was collapsed across recent memory accuracy on Day 1 was higher than on Day 14. Memory accuracy for remotely consolidated items differed between sessions in both young adults and children, showing higher remote memory accuracy on Day 1 than on Day 14. All tests used Sidak correction for multiple comparisons. Red dashed line indicates the threshold for random performance. **p* < .05; ***p* < .01; ****p* <  .001(significant difference); non-significant differences were not specifically highlighted. Error bars indicate standard error based on the underlying LME-model.

#### S2.1. Memory Strength across Time

To analyse the time-related change in the memory strength, we employed the drift diffusion modelling approach (Forstmann et al., 2016; Fudenberg et al., 2020; Ratcliff & McKoon, 2008; Wagenmakers et al., 2007a). This approach utilizes performance accuracy and reaction time. We calculated following parameters: (i) the drift rate (v), which indicates memory strength or the average rate of evidence accumulation; (ii) the boundary (a) parameter, which indicates the amount of evidence required to decide or stringency of the decision; (iii) the non-decision time (Ter), which reflects sensorimotor processing time. The analysis was based on the EZ-diffusion model (Wagenmakers et al., 2007). In this model, the parameters are estimated based on memory performance accuracy, the mean and the variance of reaction time of the correct responses. With the derived parameters, we conducted linear mixed-effect models (LME model) for memory measures using the lmer function from the lme4 package in R (Bates et al., 2015) and lmerTest (Kuznetsova et al., 2017). All LME models were calculated with maximum-likelihood estimation and Subject as the random intercept to account for between-subject variability in the derived parameters of the drift diffusion model. For that, we included the within-subject factor of *Session* (Day 0, Day 1, and Day 14) and the between-subject factor of *Group* (children and young adults) in the LME models. All main and interaction effects were False Discovery Rate adjusted for multiple comparisons.

To characterize the change in memory strength across time within and between child and adult groups, we employed the drift diffusion modelling approach (Forstmann et al., 2016; Fudenberg et al., 2020; Ratcliff & McKoon, 2008; Wagenmakers et al., 2007a) that utilizes not only performance accuracy but also reaction time in complex tasks (Criss, 2010; Lerche & Voss, 2019; Palada et al., 2016; Zhou et al., 2021) and can be applied for different developmental groups (Ratcliff et al., 2011, 2012). We calculated (i) the drift rate (v), which indicates the average rate of evidence accumulation in favour of a correct decision. Thus, the drift rate reflects accessibility of memory representations: a higher value indicates a greater probability of making a correct decision, indicating stronger memory. Conversely, lower values suggest slower accumulation of evidence, possibly indicating difficulty in processing information or a lower signal-to-noise ratio strength (Turker & Swallow, 2022). Further, we calculated also (ii) the boundary (a) parameter, which indicates the amount of evidence required to decide. Larger boundary values mean that more information is needed before deciding, leading to more accurate but slower decisions. Conversely, a smaller boundary value suggests that less information is needed, resulting in faster but potentially less accurate decisions. Lastly, (iii) the non-decision time (Ter) was calculated, reflecting the portion of response time that is not related to decision process. A low non-decision time suggests that most of response time is consumed by actual mnemonic decision process rather than peripheral processes. Conversely, a high non-decision time indicates that a large portion of response time is taken up by processes other than mnemonic decision-making.

All these parameters, namely the drift rate, boundary, and non-decision time, were calculated for children and young adults for recent (immediately retrieved), remote Day 1 and remote Day 2 memory items. For recent memory items, we aggregated the drift rates , the boundary, and non-decision time across two sessions, as there were no significant differences between sessions, as indicated by nonsignificant *Session* and *Session x Group* interactions (all p > .13). Additionally, we conducted LME model analyses for each parameter, with *Subject* as a random factor, and *Group* and *Delay* as fixed effects.

Firstly, the Linear Mixed Effects (LME) model for drift rate (v) explained a significant amount of variance R^2^ = .83, 95% CI [.83 - .88]. We observed a significant main effect of *Group*, F_(1,84)_ = 86.56, p < .001_FDR-adjusted_, w^2^ = .44, indicating a lower overall drift rate in children compared to young adults, b = -.06, t_(89)_ = -8.24, p < .001. There was also a significant *Delay* effect, F_(2,156)_ = 215.43, p < .001 _FDR-adjusted_, w^2^ = .73, showing an overall higher drift rate for recent items compared to remote Day 1 items, b = .02, t_(161)_ = 5.54, p < .001, and the drift rate was significantly higher for remote Day 1 compared to remote Day 14, b = .06, t_(165)_ = 14.64, p < .001. Additionally, there was a significant *Group x Delay* interaction, F_(2,156)_ = 28.08, p < .001 _FDR-adjusted_, w^2^ = .25. Sidak-corrected post hoc tests revealed that the slope of decrease of the drift rate from recent to remote Day 1 was more pronounced in young adults compared to children, b = -.03, t_(161)_ = -4.24, p = <.001, and the slope of decrease of the drift rate from remote Day 1 to remote Day 14 was steeper in young adults, b = -.03, t_(165)_ = -3.28, p = .008. The results show overall lower memory strength in children compared to adults, indicating less effective long-term memory consolidation in children compared to young adults already immediately after learning and extending into longer delays. Albeit adults showed higher memory strength during all delays, the decline rate was faster compared to children, indicating with this profound changes in the memory strength of initially strong memories that stronger memories tend to lose more.

Figure S2

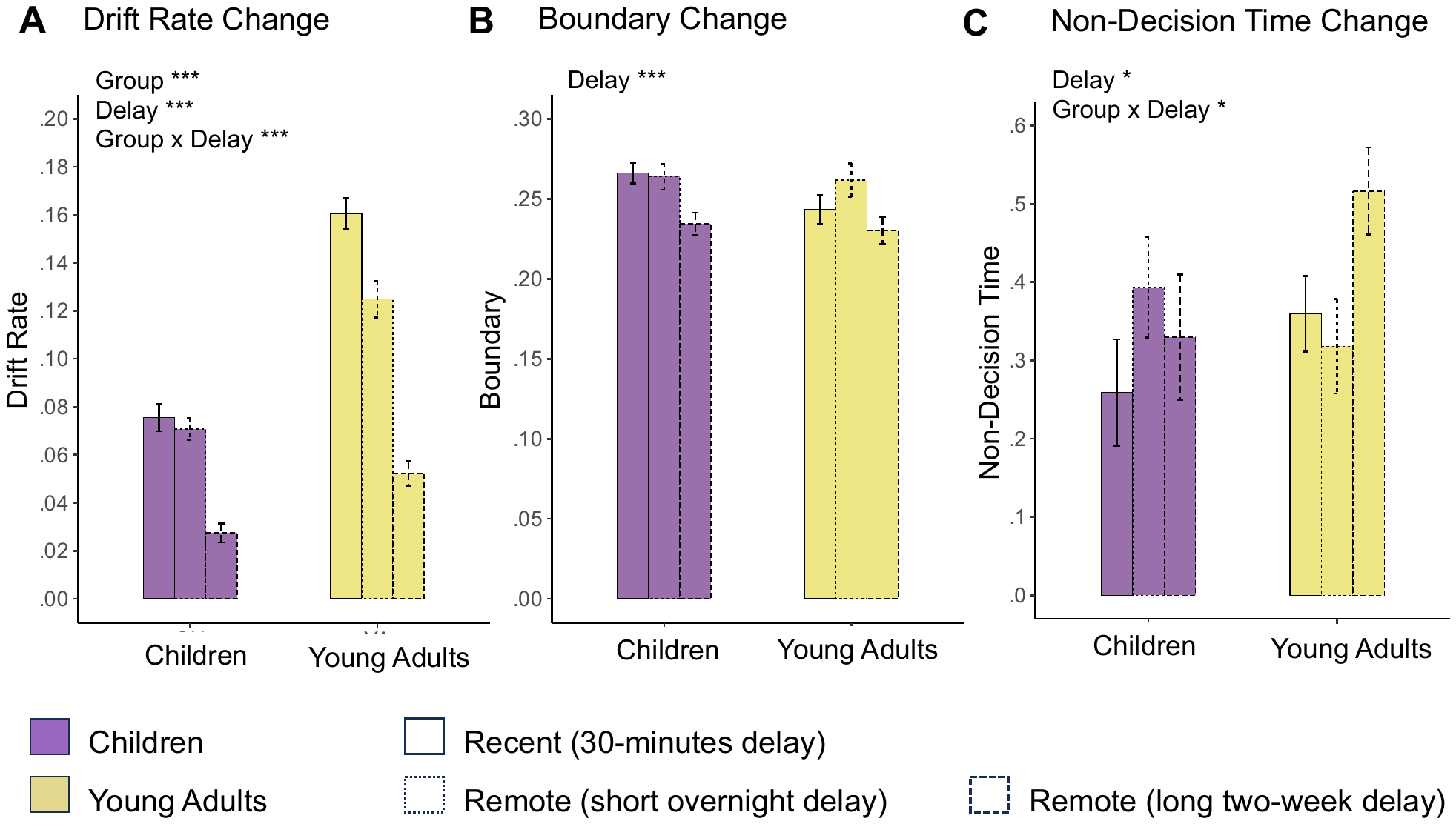

**Delay-related Change in Memory Strength as Indicated by Drift Rate, Boundary and Non-decision Time Change Within and Between Children and Young Adults.** (A) Drift Rate Change reflects the change in the memory strength or efficiency of evidence accumulation (retrieval processes) to choose a correct item location. (B) Boundary Change reflects the delay-related change in the stringency of retrieval-based decision process. (C) Non-decision time change reflects the delay-based change in sensorimotor processing during memory retrieval decision. **p* < .05; ***p* < .01; ****p* < .001(significant difference); non-significant differences were not specifically highlighted. Error bars indicate standard error based on the underlying LME-model.

Secondly, the LME model for the boundary (a) explained a significant amount of variance R2 = .43, 95% CI [.39 - .53]. It revealed a significant main effect of *Delay*, F_(2,159)_ = 11.32, p = <.001_FDR-adjusted_, w^2^ = .11. The overall boundary remained constant for recent to remote Day 1 items, b = -.008, t_(161)_ = -1.24, p = .520, but was significantly higher for remote Day 1 compared to remote Day 14 memories, b = .03, t_(167)_ = 4.57, p = <.001. Neither the *Group* effect nor the *Group x Delay* interaction was significant (all p > .227), indicating that the boundary and its change over time were similar in children and young adults. Overall, these findings indicate a slight decrease in boundary separation from a short to a long remote delay across both age groups. This decrease might suggest that participants are slightly more inclined to make mnemonic decisions with less evidence after two weeks.

Thirdly, the LME model for the non-decision time (Ter) explained a significant amount of variance R2 = .50, 95% CI [.44 - .60]. The LME revealed a significant main effect of *Delay*, F_(1,157)_ = 3.57, p = .030_FDR-adjusted_, w^2^ = .03. Sidak-adjusted post hoc tests revealed overall lower non-decision time for recent items compared to remote Day 14 items, b = -.13, t_(165)_ = -2.63, p = .028. There was no significant main effect of *Group* (p = .293), indicating similar non-decision time between children and adults. In addition, a significant *Group x Delay* interaction was observed, F_(2,157)_ = 4.32, p = .022 _FDR-adjusted_, w^2^ = .04. The Sidak-adjusted post hoc tests showed significantly higher non-decision time for remote Day 14 memories compared to remote Day 1 memories in young adult, b = .22, t_(164)_ = 3.10, p = .013. This delay-related increase in adults was significantly higher compared to children, b = .28, t_(166)_ = 2.83, p = .031. There were no other significant between or within group difference in the non-decision time (all p > .18). Overall, these findings suggest that overall increase in non-decision time over time was driven by the young adult group.

Table S2

*Statistical overview of the main and interaction effects of the linear mixed effects model for drift diffusion parameters.*

|  | **Main Effect**  **of Group** | | **Main Effect**  **of Delay** | | **Group x Delay Interaction** | |  |
| --- | --- | --- | --- | --- | --- | --- | --- |
| ***Regions of Interest*** | *F_(DF)_* | p | *F_(DF)_* | *p* | *F_(DF)_* | *p* | *R2* |
| V | 69.56(1,84) | <.001 | 215.43(2,156) | <.001 | 28.08(2,156) | <.001 | .829 |
| A | 1.42(1,85) | .293 | 11.32(2,159) | <.001 | 1.50(2,159) | .227 | .429 |
| Ter | 1.12_(1,85)_ | .293 | 3.57_(2,157)_ | .030 | 4.32_(2,157)_ | .022 | .500 |

*Notes.* Subject was included as random effect. Group (children, young adults), Delay ( recent, remote (Day 1), remote (Day 14)), and their interaction were included as fixed effect. The following reference levels where used: for Delay, recent; for Group, Children; V – drift rate; A – boundary; Ter – Non-decision Time; F – F-value; DF – degrees of freedom; p – p-value; R2 – amount of variance explained by the model (Stoffel et al., 2021). All main and interaction effects are False Discovery Rate corrected for multiple comparisons. Type III Analysis of Variance Table with Satterthwaite's method. *p < .05; ** < .01, *** < .001 (significant difference).

**S3.1. Supplementary fMRI univariate analysis**

Table S3.1

*Regions exhibiting stronger activation for remote vs. recent items in (i) young adults, (ii) children, (iii) children vs young adults, and (iv) young adults vs children on Day 1 (short delay). To capture the involved brain region better, local maxima are presented in addition to cluster maxima for the largest clusters. The preprocessing steps included global signal regression.*

| **Day 1 (Short Delay)** | | | | |  |
| --- | --- | --- | --- | --- | --- |
| **Young adults** | | | | | |
| **Region** | **x** | **y** | **x** | **Z-max** | **# voxels** |
| Left Middle Frontal Gyrus | - 44 | 2 | 40 | 6.67 | 2990 |
| Left Insula Cortex | - 34 | 22 | 2 | 6.58 |  |
| Left Inferior Frontal Gyrus, Pars Opercularis | - 44 | 6 | 34 | 6.03 |  |
| Left Lateral Occipital Cortex | - 28 | - 76 | 36 | 6.82 | 2272 |
| Left Superior Parietal Lobule | - 34 | - 50 | 44 | 5.11 |  |
| Left Fusiform Gyrus | - 44 | - 60 | - 12 | 6.7 | 1661 |
| Left Parahippocampal Gyrus | - 34 | - 34 | - 16 | 4.58 |  |
| Right Cerebellum | 30 | - 60 | - 28 | 6.03 | 1049 |
| Right Lateral Occipital Cortex | 34 | - 72 | 40 | 5.96 | 943 |
| Right Inferior Parietal Lobule | 38 | - 78 | 26 | 4.3 |  |
| Right Parahippocampal Gyrus | 32 | - 34 | - 16 | 5.29 | 718 |
| Right Inferior Temporal Gyrus | 52 | - 54 | - 10 | 5.17 |  |
| Left Superior Frontal Gyrus | - 4 | 16 | 48 | 5.04 | 405 |
| Right insular cortex | 30 | 24 | 2 | 5.25 | 279 |
| Right Middle Frontal Gyrus, Pars Triangularis | 40 | 30 | 20 | 3.61 |  |
| Right precentral Gyrus | 42 | 2 | 30 | 4.97 | 146 |
| Right Middle Frontal Gyrus, Pars Opercularis | 50 | 16 | 32 | 3.41 |  |
| Left Frontal Orbital Cortex | - 26 | 32 | - 10 | 4.51 | 123 |
| Left Cingulate Gyrus | - 4 | 2 | 28 | 4.86 | 103 |
| **Children** | | | | | |
| Right Temporal Occipital Fusiform Cortex | 26 | - 44 | - 8 | 5.1 | 658 |
| Right Parahippocampal Gyrus | 30 | - 36 | - 16 | 4.93 |  |
| Right Precuneus | 8 | - 52 | 6 | 4.79 |  |
| Left Temporal Fusiform Gyrus | - 34 | - 42 | - 12 | 5.59 | 500 |
| Left Parahippocampal Gyrus | - 18 | - 42 | - 10 | 4.91 |  |
| Left Precuneus Cortex | - 14 | - 60 | 10 | 4.47 | 160 |
| Left Lateral Occipital Cortex | - 36 | - 84 | 26 | 4.95 | 112 |
| **Children > Young Adults** | | | | | |
| Right precuneus | 4 | - 48 | 30 | 5.25 | 1051 |
| Left precuneus | - 4 | - 48 | 40 | 4.68 |  |
| Right Superior Parietal Lobule | 12 | - 32 | 50 | 4.99 | 203 |
| Right Parietal Operculum Cortex | 54 | - 30 | 24 | 3.32 | 149 |
| **Young Adults > Children** | | | | | |
| Left Precentral Gyrus, Middle Frontal Gyrus | - 44 | 2 | 40 | 4.8 | 501 |
| Left Inferior Frontal Gyrus | - 54 | 14 | 10 | 3.39 |  |
| Left Frontal Operculum Cortex | - 34 | 22 | 2 | 5.48 | 260 |
| Right Cerebellum | 12 | - 76 | - 20 | 4.7 | 141 |
| Left Medial Frontal Gyrus | - 2 | 16 | 48 | 4.2 | 118 |
| Left/Right Insular Cortex | 32 | 22 | 2 | 4.66 | 113 |
| Left/Right Lateral Occipital Cortex | - 26 | - 74 | 36 | 4.5 | 107 |

Table S4.1

*Regions exhibiting stronger activation for remote vs. recent items in (i) young adults, (ii) children, (iii) children vs young adults, and (iv) young adults vs children on Day 14 (long delay). To capture the involved brain region better, local maxima are presented in addition to cluster maxima for the largest clusters. The preprocessing steps included global signal regression.*

| **Day 14 (Long Delay)** | | | | | |
| --- | --- | --- | --- | --- | --- |
| **Young Adults** | | | | | |
| **Region** | **x** | **y** | **x** | **Z-max** | **# voxels** |
| Left/Right Occipital Fusiform Gyrus | - 46 | - 58 | - 16 | 7.62 | 19227 |
| Left Lateral Occipital Cortex | - 30 | - 60 | - 14 | 7.25 |  |
| Left Middle Frontal Gyrus, Pars Opercularis, |  |  |  | 7.17 | 2890 |
| Left Superior Frontal Gyrus | - 6 | 12 | 56 | 6.78 |  |
| Right Inferior Frontal Gyrus, Pars Opercularis, Pars Trinagularis | 46 | 12 | 28 | 6 | 691 |
| Left Insular Cortex | - 32 | 22 | 2 | 6.7 | 501 |
| Left Caudate | - 10 | 4 | 10 | 5.58 | 456 |
| Right Frontal Orbital Cortex | 34 | 28 | 0 | 6.11 | 298 |
| Right Cerebellum | 16 | - 44 | - 46 | 4.97 | 250 |
| Right Caudate | 8 | 12 | 2 | 5.27 | 215 |
| Left Cerebellum | - 34 | - 68 | - 54 | 6.1 | 211 |
| **Children** | | | | | |
| Left Temporal Fusiform Gyrus | - 34 | - 26 | - 24 | 4.91 | 580 |
| Left anterior Parahippocampal Gyrus, Hippocampus | - 36 | - 18 | - 24 | 4.4 |  |
| Left Lateral Occipital Cortex | - 48 | - 58 | - 16 | 4.25 |  |
| Right Temporal Occipital Fusiform Cortex | 40 | - 54 | - 18 | 4.34 | 448 |
| Right Lateral Occipital Cortex | 50 | - 70 | - 12 | 4.2 |  |
| **Children > Young Adults** | | | | | |
| Right/Left angular gyrus | 62 | - 40 | 44 | 4.8 | 847 |
| Right/Left Lateral Occipital Cortex | 46 | - 66 | 48 | 4.44 |  |
| Right Superior Frontal Gyrus | 20 | 30 | 58 | 4.58 | 640 |
| Right/Left Superior Temporal Gyrus |  |  |  | 4.73 | 493 |
| Right Precuneus | 8 | - 52 | 30 | 4.51 | 332 |
| Right Medial Frontal Cortex | 8 | 50 | - 2 | 4.35 | 287 |
| Right Middle Temporal Gyrus | 66 | - 18 | - 20 | 4.17 | 203 |
| Left Middle Frontal Gyrus | - 20 | 36 | 38 | 4.31 | 154 |
| Left Cingulate Gyrus | - 14 | - 50 | 30 | 4.36 | 138 |
| **Young Adults > Children** | | | | | |
| Right/Left Cerebellum | 14 | - 72 | - 22 | 5.77 | 3162 |
| Left Occipital Fusiform Gyrus | - 20 | - 90 | - 14 | 5.22 | 1229 |
| Left Lateral Occipital Cortex | - 30 | - 80 | 36 | 5.62 | 620 |
| Left Middle Frontal Gyrus, Inferior Frontal Gyrus | - 44 | 12 | 30 | 4.8 | 387 |
| Right Precuneus | 18 | - 58 | 20 | 4.39 | 205 |
| Left Superior Frontal Gyrus | - 6 | 12 | 56 | 5.12 | 165 |
| Left Posterior Parahippocampal Gyrus, Hippocampus | - 28 | - 32 | - 18 | 3.9 | 96 |

Table S5.1

*Regions exhibiting stronger activation for remote vs. recent items that decreases over time (i) in young adults stronger than in children (ii) children stronger than in adults; that increases over time (iii) in young adults stronger than in children, and (iv) in children stronger than in young adults. To capture the involved brain region better, local maxima are presented in addition to cluster maxima for the largest clusters. The preprocessing steps included global signal regression.*

| **Decrease Across Time** | | | | | |
| --- | --- | --- | --- | --- | --- |
| **Young Adults > Children** | | | | | |
| **Region** | **x** | **y** | **x** | **Z-max** | **# voxels** |
| Right Superior Parietal Lobule, Agular Gyrus | 42 | - 50 | 58 | 3.69 | 946 |
| Right Middle Frontal Gyrus | 42 | 56 | 2 | 4.16 | 546 |
| Left Middle Frontal Gyrus | - 38 | 24 | 48 | 3.9 | 379 |
| Right Superior Frontal Gyrus | 8 | 48 | 30 | 3.44 | 329 |
| **Children > Adults** | | | | | |
| Left Lateral Occipital Cortex | - 32 | - 88 | 6 | 4.81 | 4474 |
| Left Hippocampus, Posterior Parahippocampal Gyrus | - 30 | - 30 | - 6 | 4.09 |  |
| Right Lateral Occipital Cortex, Occipital Fusiform Gyrus, Lingual Gyrus | 30 | - 86 | 4 | 4.73 | 1717 |
| **Increase Over Time** | | | | | |
| **Young Adults > Children** | | | | | |
| Left Lateral Occipital Cortex | - 32 | - 88 | 6 | 4.81 | 4474 |
| Left Hippocampus | - 30 | - 30 | - 6 | 4.09 |  |
| Left Lingual gyrus | - 10 | - 56 | - 6 | 4.04 |  |
| Right Lateral Occipital Cortex, Occipital Fusiform Gyrus, Precuneus | - 30 | 86 | 4 | 4.73 | 1717 |
| **Children > Young Adults** | | | | | |
| Right Superior Parietal Lobule, Angular Gyrus | 42 | - 50 | 58 | 3.69 | 946 |
| Right Middle Frontal Gyrus | 42 | 56 | 2 | 4.16 | 546 |
| Left Middle Frontal Gyrus, Superior Frontal Gyrus | - 38 | 24 | 48 | 3.9 | 379 |
| Right Superior Frontal Gyrus, Paracingulate Gyrus | 8 | 48 | 30 | 3.44 | 329 |

Table S3.2

*Regions exhibiting stronger activation for remote vs. recent items in (i) young adults, (ii) children, (iii) children vs young adults, and (iv) young adults vs children on Day 1 (short delay). To capture the involved brain region better, local maxima are presented in addition to cluster maxima for the largest clusters. The preprocessing steps did not include global signal regression.*

| **Day 1 (Short Delay)** | | | | |  |
| --- | --- | --- | --- | --- | --- |
| **(i)Young adults** | | | | | |
| **Region** | **x** | **y** | **x** | **Z-max** | **# voxels** |
| Left Lateral Occipital Cortex | - 32 | -78 | 32 | 6.07 | 1598 |
| Left Inferior Temporal Gyrus, Occipital Fusiform Gyrus | - 44 | - 60 | -12 | 6.01 | 10855 |
| Left Posterior Temporal Fusiform Gyrus | - 30 | - 42 | -14 | 4.94 |  |
| Left Cerebral White Matter | - 34 | - 54 | -4 | 4.46 | 1661 |
| Left Parahippocampal Gyrus, Posterior | - 36 | - 22 | -26 | 4.42 |  |
| Precentral Gyrus, Middle Frontal Gyrus | -44 | 0 | 40 | 6.13 | 851 |
| Inferior Frontal Gyrus, Pars Opercularis | -50 | 10 | 18 | 3.98 |  |
| Right Lateral Occipital Cortex | 28 | -68 | 48 | 5.19 | 533 |
| Left Superior/Middle Frontal Gyrus | -24 | 0 | 50 | 5.6 | 345 |
| Left Inferior Frontal Gyrus, Pars Triangularis | -54 | 34 | 4 | 3.8 | 273 |
| Right Posterior Parahippocampal Gyrus, Posterior | 22 | -34 | -12 | 4.47 | 256 |
| Cerebellum | 10 | -76 | -22 | 4.2 | 247 |
| Right Middle Inferior Temporal Gyrus | 52 | -54 | -12 | 4.41 | 195 |
| Right Insular Cortex | 30 | 22 | 2 | 4.98 | 172 |
| Right Middle Frontal Gyrus | 34 | 32 | 16 | 3.36 | 171 |
| **(ii)Children** | | | | | |
| Left Temporal Fusiform Cortex, | -36 | -40 | -12 | 4.76 | 513 |
| Left Parahippocampal Gyrus, Posterior | -16 | -42 | -10 | 4.71 |  |
| Right Temporal Fusiform Gyrus | 40 | -30 | -18 | 4.77 | 392 |
| Right Parahippocampal Gyrus, Posterior | 26 | -38 | -10 | 4.36 |  |
| **(iii)Children > Young Adults** | | | | | |
| Right Pariental Cortex, Precuneus | 4 | -48 | 30 | 5.16 | 1031 |
| Precuneus | -4 | -48 | 38 | 4.23 |  |
| **(iv)Young Adults > Children** | | | | | |
| Left Middle Frontal Gyrus | -44 | 2 | 40 | 4.02 | 204 |
| Left Inferior Frontal Gyrus, Pars Opercularis | -52 | 8 | 32 | 3.86 |  |
| Left Frontal Orbital Cortex, Operculum Cortex | -38 | 22 | -4 | 4.04 | 151 |
| Left Frontal Pole | -48 | 44 | 4 | 4.08 | 127 |

Table S4.2

*Regions exhibiting stronger activation for remote vs. recent items in (i) young adults, (ii) children, (iii) children vs young adults, and (iv) young adults vs children on Day 14 (long delay). To capture the involved brain region better, local maxima are presented in addition to cluster maxima for the largest clusters. The preprocessing steps did not include global signal regression.*

| **Day 14 (Long Delay)** | | | | | |
| --- | --- | --- | --- | --- | --- |
| **(i)Young Adults** | | | | | |
| **Region** | **x** | **y** | **x** | **Z-max** | **# voxels** |
| Right Inferior Temporal Gyrus | 50 | -58 | -14 | 7.6 | 22203 |
| Left Lateral Occipital Cortex | -34 | -84 | 22 | 7.47 |  |
| Right Temporal Fusiform Cortex | 32 | -38 | -18 | 7.47 |  |
| Left Inferior Frontal Gyrus, Pars Opercularis | -42 | 12 | 28 | 7.1 | 2319 |
| Middle Frontal Gyrus | -50 | 22 | 30 | 6.67 |  |
| Right Inferior Frontal Gyrus | 42 | 28 | 22 | 4.54 | 1084 |
| Right Middle Frontal Gyrus | 44 | 32 | 20 | 4.48 |  |
| Caudate | -12 | 12 | 4 | 5.24 | 1101 |
| Left Frontal Orbital Cortex | -36 | 34 | -10 | 5.09 | 517 |
| Cerebellum | 22 | -34 | -42 | 6.95 | 369 |
| **(ii)Children** | | | | | |
| Left Temporal Fusiform Cortex | -34 | -26 | -24 | 4.78 | 541 |
| Left Parahippocampal Gyrus, Anterior Devision | -36 | -16 | -24 | 4.41 |  |
| Left Lateral Occipital Cortex | 50 | -70 | -10 | 4.64 | 509 |
| **(iii)Children > Young Adults** | | | | | |
| Right Angular Gyrus, Parietal Lobe | 60 | -52 | 38 | 4.93 | 672 |
| Right Lateral Occipital Cortex | 46 | -66 | 48 | 4.4 |  |
| Right Cingulate Gyrus | 8 | -50 | 30 | 4.41 | 310 |
| Right Precuneus Cortex | 10 | -54 | 38 | 4.36 |  |
| Right Parietal Operculum Cortex | 54 | -26 | 22 | 3.87 | 241 |
| Right Frontal Medial Cortex | 10 | 50 | -2 | 4.32 | 141 |
| Left Precuneus Cortex | -6 | -50 | 40 | 4.45 | 131 |
| Left Insula Cortex | -32 | 8 | 12 | 3.97 | 109 |
| **(iv)Young Adults > Children** | | | | | |
| Right Cerebellum | 14 | -72 | -24 | 5.05 | 1471 |
| Left cerebellum | -30 | -80 | 36 | 5.1 | 514 |
| Left Lateral Occipital Cortex | -34 | -84 | 22 | 3.65 |  |
| Left Occipital Pole | -18 | -94 | -14 | 4.4 | 492 |
| Right Lateral Occipital Cortex | 36 | -76 | 34 | 4.57 | 366 |
| Left Middle Frontal Gyrus | -44 | 12 | 30 | 4.36 | 235 |
| Cerebellum | -32 | -68 | -54 | 5.25 | 177 |
| Left Precuneus Cortex | -16 | -62 | 18 | 4.86 | 108 |

Table S5.2

*Regions exhibiting stronger activation for remote vs. recent items that decreases over time (i) in young adults stronger than in children, (ii) children stronger than in adults, in (iii) young adult and (iv) children; that increases over time (iii) in young adults stronger than in children, and (iv) in children stronger than in young adults, in (iii) young adult and (iv) children. To capture the involved brain region better, local maxima are presented in addition to cluster maxima for the largest clusters. The preprocessing steps did not include global signal regression.*

| **Decrease Across Time** | | | | | |
| --- | --- | --- | --- | --- | --- |
| **(ii)Children > Adults** | | | | | |
| **Region** | **x** | **y** | **x** | **Z-max** | **# voxels** |
| Right Occipital Pole | 18 | -92 | 2 | 4.48 | 1195 |
| Left Lateral Occipital Cortex | -32 | -88 | 6 | 4.39 | 1141 |
| Right Precuneus Cortex | 10 | -52 | 10 | 3.66 | 588 |
| **(iii)Adults** | | | | | |
| Right Angular Gyrus | 60 | -50 | 40 | 4.02 | 1533 |
| Right Lateral Occipital Cortex | 52 | -58 | 46 | 3.5 |  |
| Right Middle Temporal Gyrus | 70 | -22 | -18 | 4.39 | 641 |
| Right Frontal Pole | 44 | 56 | -6 | 4.89 | 543 |
| Left Lateral Occipital Cortex | -46 | -62 | 38 | 4.1 | 378 |
| **Increase Over Time** | | | | | |
| **(i)Young Adults > Children** | | | | | |
| Right Occipital Pole | 18 | -92 | -14 | 4.48 | 1195 |
| Left Lateral Occipital Cortex | -32 | -88 | 6 | 4.39 | 1141 |
| Right Precuneus Cortex | 10 | -52 | 10 | 3.66 | 588 |
| **(iii)Young Adults** | | | | | |
| Left Lateral Occipital Cortex | -42 | -84 | -10 | 6.17 | 16779 |
| Right Lateral Occipital Cortex | 36 | -88 | 8 | 5.97 |  |
| Left Caudate, Thalamus, |  |  |  | 4.25 | 1455 |
| Lfer Insular Cortex, Inferior Frontal Gyrus, Triangularis, Frontal Orbital Cortex | -30 | 28 | 6 | 3.95 |  |
| Left Middle Frontal Gyrus | -38 | 18 | 32 | 4.67 | 759 |
| Left Inferior Frontal Gyrus, Pars Opercularis | -44 | 14 | 28 | 4.44 |  |
| Right Inferior Frontal Gyrus, Pars Opercularis, Pars Triangularis, | 46 | 12 | 28 | 3.6 | 468 |
| Right Middle Frontal Gyrus | 46 | 24 | 28 | 3.13 |  |
| **(iv)Children** | | | | | |
| Left Insular Cortex, Frontal Operculum Cortex | -32 | 12 | 12 | 4.14 | 1262 |
| Left Frontal Orbital Cortex | -40 | 22 | -4 | 3.54 |  |
| Left Inferior Frontal Gyrus, Pars Opercularis | -32 | 22 | 22 | 3.47 |  |

Table S6

Two-sided permutation t-tests were conducted to assess whether the mean signal difference of the contrast **remote > recent** significantly differed from zero for each combination of ROI, session, and group. For each subset, the mean signal difference value, t-statistic, and unadjusted and FDR-adjusted p-values are reported. P-values were corrected for multiple comparisons using the False Discovery Rate (FDR) method.

| **ROI** | **Session** | **Group** | **Mean value** | **t statistic** | **p-value** | **Adjusted**  **p-value** |
| --- | --- | --- | --- | --- | --- | --- |
| **Children – Short Delay** | | | | | | |
| Medial Prefrontal Cortex | Day 1 | Children | .056 | 2.007 | .051 | .102 |
| Cerebellum | Day 1 | Children | **.047** | **2.937** | **.005** | **.017** |
| Retrosplenial Cortex | Day 1 | Children | .046 | 2.437 | .019 | .054 |
| Precuneus | Day 1 | Children | .019 | .920 | .362 | .451 |
| Hippocampus Anterior | Day 1 | Children | .027 | 2.034 | .048 | .102 |
| Hippocampus Posterior | Day 1 | Children | .018 | 1.301 | .200 | .307 |
| Parahippocampus Anterior | Day 1 | Children | .007 | .456 | .651 | .723 |
| Parahippocampus Posterior | Day 1 | Children | **.072** | **3.719** | **<.001** | **.002** |
| Ventrolateral Prefrontal Cortex | Day 1 | Children | .018 | .881 | .383 | .451 |
| Lateral Occipital Cortex | Day 1 | Children | .018 | 1.009 | .318 | .433 |

| **Young Adults – Short Delay** | | | | | | |
| --- | --- | --- | --- | --- | --- | --- |
| Medial Prefrontal Cortex | Day 1 | Young Adults | .003 | .121 | .902 | .927 |
| Cerebellum | Day 1 | Young Adults | .028 | 2.396 | .022 | .058 |
| Retrosplenial Cortex | Day 1 | Young Adults | -.010 | -.882 | .383 | .451 |
| Precuneus | Day 1 | Young Adults | **-.062** | **-4.196** | **<.001** | **.001** |
| Hippocampus Anterior | Day 1 | Young Adults | .028 | 2.169 | .036 | .086 |
| Hippocampus Posterior | Day 1 | Young Adults | .012 | 1.526 | .068 | .135 |
| Parahippocampus Anterior | Day 1 | Young Adults | .019 | 1.539 | .326 | .433 |
| Parahippocampus Posterior | Day 1 | Young Adults | **.063** | **5.431** | **<.001** | **<.001** |
| Ventrolateral Prefrontal Cortex | Day 1 | Young Adults | **.129** | **6.455** | **<.001** | **<.001** |
| Lateral Occipital Cortex | Day 1 | Young Adults | .022 | 1.387 | .173 | .277 |

| **Children – Long Delay** | | | | | | |
| --- | --- | --- | --- | --- | --- | --- |
| Medial Prefrontal Cortex | Day 14 | Children | **.103** | **3.252** | **.002** | **.009** |
| Cerebellum | Day 14 | Children | .046 | 1.586 | .120 | .229 |
| Retrosplenial Cortex | Day 14 | Children | -.031 | -1.393 | .171 | .277 |
| Precuneus | Day 14 | Children | **-.063** | **-2.609** | **.013** | **.039** |
| Hippocampus Anterior | Day 14 | Children | .009 | .376 | .709 | .758 |
| Hippocampus Posterior | Day 14 | Children | .007 | .361 | .720 | .758 |
| Parahippocampus Anterior | Day 14 | Children | .027 | .995 | .282 | .402 |
| Parahippocampus Posterior | Day 14 | Children | .047 | 2.023 | .050 | .102 |
| Ventrolateral Prefrontal Cortex | Day 14 | Children | **.073** | **3.055** | **.004** | **.014** |
| Lateral Occipital Cortex | Day 14 | Children | .049 | 2.281 | .028 | .069 |

| **Young Adults – Long Delay** | | | | | | |
| --- | --- | --- | --- | --- | --- | --- |
| Medial Prefrontal Cortex | Day 14 | Young Adults | .002 | .071 | .944 | .944 |
| Cerebellum | Day 14 | Young Adults | **.113** | **6.216** | **<.001** | **<.001** |
| Retrosplenial Cortex | Day 14 | Young Adults | .021 | 1.270 | .213 | .315 |
| Precuneus | Day 14 | Young Adults | **-.089** | **-4.046** | **<.001** | **.001** |
| Hippocampus Anterior | Day 14 | Young Adults | .014 | .977 | .336 | .433 |
| Hippocampus Posterior | Day 14 | Young Adults | .005 | .580 | .566 | .646 |
| Parahippocampus Anterior | Day 14 | Young Adults | .015 | 1.095 | .141 | .235 |
| Parahippocampus Posterior | Day 14 | Young Adults | **.147** | **8.058** | **<.001** | **<.001** |
| Ventrolateral Prefrontal Cortex | Day 14 | Young Adults | **.242** | **9.325** | **<.001** | **<.001** |
| Lateral Occipital Cortex | Day 14 | Young Adults | **.148** | **6.287** | **<.001** | **<.001** |

*Notes*. ROI – region of interest; p – p-value; *p < .05; ** < .01, *** < .001 (significant difference).

Table S7

*Test of neural activation during object presentation separately for recent and remote memories for significance (higher than zero).*

|  | Recent | | | | | Remote | | |
| --- | --- | --- | --- | --- | --- | --- | --- | --- |
|  | Young Adults | | | | | | | |
| ROI | *Day* | *mean* | *T test* | *p_(FDRadj)_* | *mean* | | *T test* | *p_(FDRadj)_* |
| Hippocampus Anterior | Day 1 | .054 | 3.76 | **<.001** | .083 | 6.42 | | **<.001** |
|  | Day 14 | .072 | 6.25 | **<.001** | .089 | 6.95 | | **<.001** |
| Hippocampus Posterior | Day 1 | .056 | 5.79 | **<.001** | .069 | 6.91 | | **<.001** |
|  | Day 14 | .063 | 7.71 | **<.001** | .068 | 6.66 | | **<.001** |
| Parahippocampal Gyrus Anterior | Day 1 | .025 | 1.98 | **.031** | .044 | 3.45 | | **.001** |
|  | Day 14 | .038 | 2.59 | **.010** | .054 | 4.31 | | **<.001** |
| Retrosplenial Cortex | Day1 | .120 | 6.75 | **<.001** | .113 | 5.64 | | **<.001** |
|  | Day14 | .079 | 5.39 | **<.001** | .108 | 7.24 | | **<.001** |
| Precuneus | Day1 | .099 | 5.37 | **<.001** | .034 | 1.808 | | **.041** |
|  | Day14 | .150 | 8.48 | **<.001** | .057 | 3.79 | | **<.001** |
|  | Children | | | | | | | |
|  |  | *mean* | *T test* | *p_(FDRadj)_* | *mean* | *T Test* | | *p_(FDRadj)_* |
| Hippocampus Anterior | Day 1 | .043 | 2.51 | **.011** | .076 | 4.47 | | **<.001** |
|  | Day 14 | .080 | 4.09 | **<.001** | .092 | 3.37 | | **.001** |
| Hippocampus Posterior | Day 1 | .017 | 1.09 | .141 | **.037** | **2.405** | | **.013** |
|  | Day 14 | .035 | 2.28 | **.016** | .048 | 2.45 | | **.013** |
| Parahippocampal Gyrus Anterior | Day 1 | .058 | 3.69 | **.001** | .064 | 3.81 | | **.000** |
|  | Day 14 | .070 | 2.64 | **.009** | .098 | 3.14 | | **.002** |
| Retrosplenial Cortex | Day1 | .042 | 1.97 | **.031** | .089 | 4.28 | | **<.001** |
|  | Day14 | .090 | 4.59 | **<.001** | .055 | 2.37 | | **.014** |
| Precuneus | Day1 | .036 | 1.63 | .056 | .052 | 1.31 | | **.013** |
|  | Day14 | .130 | 6.72 | **<.001** | .053 | 1.88 | | **.037** |

*Notes*.To test for significance we used one-sample permutation t-test for more robust calculations with Monte-Carlo permutation percentile confidence interval. All p-values for False Discovery Rate (FDR) corrected for multiple comparisons. ROI – region of interest; p – p-value; FDRadj – False Discovery Rate adjustment; *p < .05; ** < .01, *** < .001 (significant difference).

**Table S8**

*Statistical overview of the main and interaction effects of the linear mixed effects models recent and remote univariate results of correctly recognized items.*

|  | **Hippocampus Anterior** | | **Hippocampus Posterior** | | **Parahippocampal Cortex Anterior** | |
| --- | --- | --- | --- | --- | --- | --- |
|  | *F_(DF)_* | *p-value* | *F_(DF)_* | *p-value* | *F_(DF)_* | *p-value* |
| Delay: (Recent > Remote) | 3.62_(231)_ | .058 | 2.17_(234)_ | .142 | 1.70_(227)_ | .194 |
| Group: (Adults > Children) | .01_(80)_ | .912(.912) | **5.62_(85)_** | **.020(.060)** | 4.77_(76)_ | .030(.060) |
| Session: (Day1> Day14) | 2.65_(253)_ | .106 | 1.24_(253)_ | .267 | 1.48_(250)_ | .225 |
| Delay x Group | .001_(253)_ | .977 | .155_(234)_ | .694 | .000_(227)_ | .994 |
| Delay x Session | .462_(253)_ | .490 | .172_(234)_ | .679 | .110_(227)_ | .740 |
| Group x Session | .486_(253)_ | .486 | .670_(253)_ | .414 | .206_(250)_ | .650 |
| Delay x Group x Session | .022_(231)_ | .880 | .000_(234)_ | .990 | .215_(222)_ | .643 |
|  | **Medial Prefrontal Cortex** | | **Precuneus** | | **Retrosplenial Cortex** | |
| Delay: (Recent > Remote) | 6.82_(235)_ | .009_()_ | 15.14_(235)_ | .0001_()_ | .49_(229)_ | .484 |
| Group: (Adults > Children) | 7.86_(85)_ | .006_()_ | .959_(84)_ | **.303_(.)_** | 3.22_(80)_ | **.076(.101)** |
| Session: (Day1> Day14) | 17.74_(256)_ | <.001 | 9.49_(256)_ | .002 | .306_(247)_ | .581 |
| Delay x Group | 6.89 _(235)_ | .009 | 2.87_(235)_ | .091 | .034_(229)_ | .853 |
| Delay x Session | .873_(235)_ | .351_()_ | 4.74_(235)_ | .030_(.)_ | .879_(229)_ | .340 |
| Group x Session | 6.75_(256)_ | .009 | .196_(256)_ | .657 | 2.07_(248)_ | .152 |
| Delay x Group x Session | .477_(235)_ | .490_()_ | 1.29_(233)_ | .256_(.)_ | 5.756_(229)_ | .017 |
|  | **Ventrolateral Prefrontal Cortex** | | **Cerebellum** | | **Parahippocampal Cortex Posterior** | |
| Delay: (Recent > Remote) | 80.52_(231)_ | <.001_()_ | 28.66_(232)_ | <.001_(<.001)_ | 50.25_(233)_ | <.001_(<.001)_ |
| Group: (Adults > Children) | 5.43_(83)_ | .022_()_ | .808_(81)_ | .371_()_ | 1.55_(84)_ | .217_()_ |
| Session: (Day1> Day14) | 11.52_(248)_ | <.001 | .002_(254)_ | .965 | 34.79_(251)_ | <.001 |
| Delay x Group | 27.45_(231)_ | <.001 | .783_(232)_ | .377 | 2.59_(233)_ | .109 |
| Delay x Session | 10.82_(231)_ | .001_()_ | 4.29_(232)_ | .039_()_ | 1.87_(233)_ | .172_()_ |
| Group x Session | .004_(248)_ | .948 | 1.24_(248)_ | .266 | .671_(251)_ | .414 |
| Delay x Group x Session | 1.275_(231)_ | .260_()_ | 3.99_(232)_ | .047_()_ | 4.218_(233)_ | .041_()_ |
|  | **Lateral Occipital Cortex** | |  | | | |
| Delay: (Recent > Remote) | 21.44_(233)_ | <.001_(<.001)_ |  |  |  |  |
| Group: (Adults > Children) | 30.07_(86)_ | <.001 |  |  |  |  |
| Session: (Day1> Day14) | 46.66_(246)_ | <.001 |  |  |  |  |
| Delay x Group | 3.28_(233)_ | .071 |  |  |  |  |
| Delay x Session | 7.46_(233)_ | .007_()_ |  |  |  |  |
| Group x Session | 1.18_(246)_ | .278 |  |  |  |  |
| Delay x Group x Session | 1.29_(233)_ | .257_(.)_ |  |  |  |  |

*Notes.* Subject was included as a random effect. Group (children, young adults), Delay ( recent, remote), Session (Day1, Day 14), and their interaction were included as fixed effect. The following reference levels where used: for Delay, recent; for Group, Children; F – F-value; DF – degrees of freedom; p – p-value; Type III Analysis of Variance Table with Satterthwaite's method. *p < .05; ** <.01, ***<.001 (significant difference).

Figure S3.

| 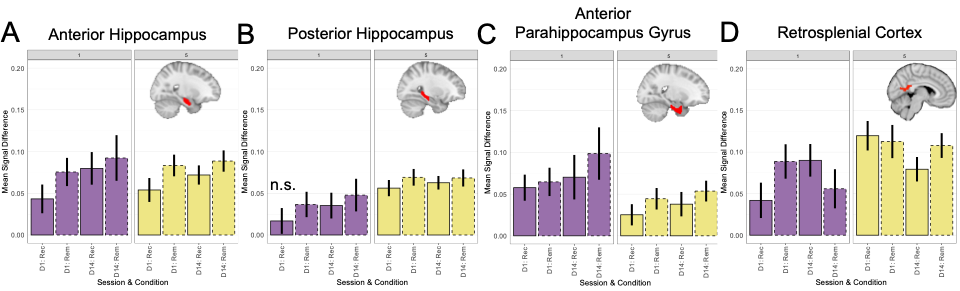 |
| --- |

**Mean Blood Oxygen Level-Dependent (BOLD) Signal Intensity**

The figure presents the mean blood oxygen level-dependent (BOLD) signal intensity for recent and remote memories on Day 1 and Day 14 in children and adults in **(A)** anterior hippocampus; (**B**) posterior hippocampus; **(C)** anterior parahippocampal gyrus. *Note:* Bars represent the average BOLD signal intensity. The colour indicated the age groups: purple for children and khaki yellow for young adults. Solid-lined bars represent recent data from Day 1 or Day 14, while dashed-lined bars depict remote data from Day1 and Day 14. Error bars indicate standard error of the mean.

Table S9

*Full statistical overview of LME model for univariate analysis (includes only ROI that show above or below zero upregulation).*

|  | **Main Effect**  **of Group** | | **Main Effect**  **of Session** | | **Group x Session Interaction** | |  |
| --- | --- | --- | --- | --- | --- | --- | --- |
| ***Regions of Interest*** | *F_(DF)_* | *p* | *F_(DF)_* | *p* | *F_(DF)_* | p | *R2* |
| Parahippocampal Gyrus Posterior | 2.97_(1,83)_ | .088_(.106)_ | 2.48_(1,100)_ | .118_(.142)_ | 9.54_(1,83)_ | **.002_(.012)_** | .200⊥ |
| Medial Prefrontal Cortex | 7.61_(1,86)_ | **.007_(.014)_** | .42_(1,99)_ | .517_(.517)_ | 1.16_(1,83)_ | .284_(.284)_ | .369⊥ |
| Ventrolateral Prefrontal Cortex | 31.35_(1,82)_ | **<.001_(<.001)_** | 10.68_(1,99)_ | **.001_(.003)_** | 1.61_(1,83)_ | .207_(.248)_ | .309⊥ |
| Cerebellum | 1.54_(1,161)_ | .215_(.215)_ | 4.67_(1,161)_ | .036_(.054_**_)_** | 7.68_(1,161)_ | **.006_(.018)_** | .100∩ |
| Precuneus | 5.09_(1,161)_ | **.025_(.037)_** | 6.50_(1,161)_ | **.011_(.022)_** | 1.61_(1,161)_ | .205_(.248)_ | .099∩ |
| Lateral Occipital Cortex | 9.12_(1,82)_ | **.003_(.009)_** | 16.76_(1,97)_ | **<.001_(<.001)_** | 6.42_(1,81)_ | **.013_(.026)_** | .324⊥ |

|  | **Main Effect**  **of Sex** | | **Main Effect**  **of Handedness** | | **Main Effect**  **of IQ** | | **Main Effect**  **Of Reaction Time** | |
| --- | --- | --- | --- | --- | --- | --- | --- | --- |
| ***Regions of Interest*** | *F_(DF)_* | *p* | *F_(DF)_* | *p* | *F_(DF)_* | p | *F_(DF)_* | *p* |
| Parahippocampal Gyrus Posterior | 1.26_(1,84)_ | .263 | .03_(1,93)_ | .962 | 1.28_1,84)_ | **.**259 | .09_(1,155)_ | .764 |
| Medial Prefrontal Cortex | .62_(1,87)_ | .430 | .50_(1,95)_ | .607 | 5.16_(1,87)_ | **.024** | .22_(1,160)_ | .635 |
| Ventrolateral Prefrontal Cortex | 1.11_(1,83)_ | .294 | .71_(1,92)_ | .494 | .09_(1,83)_ | .764 | .20_(1,154)_ | .654 |
| Cerebellum | 3.15_(1,161)_ | .077 | .21_(1,161)_ | .806 | .781_(1,161)_ | .378 | .11_(1,161)_ | .741 |
| Precuneus | .35_(1,161)_ | .553 | .20_(1,161)_ | .817 | .08_(1,161)_ | .776 | .137_(1,161)_ | .712 |
| Lateral Occipital Cortex | .10_(1,83)_ | .752 | .76_(1,92)_ | .468 | 3.04_(1,83)_ | .084 | .005_(1,159)_ | .944 |

*Notes.* *Notes.* Subject was included as random effect. Group (children, young adults), Session (Day 1 remote > recent, Day 14 remote > recent), and their interaction were included as fixed effect. The following reference levels where used: for Session – Day 1; for Group – Children; F – F-value; DF – degrees of freedom; p – p-value; FDR_adj – False Discovery Rate adjusted; R2 – amount of variance explained by the model (∩- marginal; ⊥ - conditional). Type III Analysis of Variance Table with Satterthwaite’s method. *p < .05; ** < .01, *** < .001 (significant difference). All p-values of main and interactions effects were FDR-adjusted for multiple comparisons.

Table S9.1

*Full statistical overview of LME model for univariate analysis (sub-sampled to those participants who reached accuracy criteria after 2 encoding loops).*

|  | **Main Effect**  **of Group** | | | **Main Effect**  **of Session** | | **Group x Session Interaction** | |  |
| --- | --- | --- | --- | --- | --- | --- | --- | --- |
| ***Regions of Interest*** | *F_(DF)_* | | *p* | *F_(DF)_* | *p* | *F_(DF)_* | p | *R2* |
| Hippocampus Anterior | .02_(1,125)_ | .894 | | .931_(1,125)_ | .336_(.707)_ | .701_(1,125)_ | .404_(.494)_ | .042⊥ |
| Hippocampus Posterior | 1.62_(1,125)_ | | .206_(_ | 3.51_(1,125)_ | .063_(.939)_ | 1.73_(1,125)_ | .191_(.607)_ | .141⊥ |
| Parahippocampal Gyrus Anterior | .147_(1,125)_ | | .702 | .496_(1,125)_ | .482_(.707)_ | .110_(1,125)_ | .740_(.680)_ | .051∩ |
| Parahippocampal Gyrus Posterior | 5.22_(1,61)_ | | .021 | .440_(1,71)_ | .509_(.330)_ | 22.05_(1,58)_ | <.001_(.095)_ | .482⊥ |
| Medial Prefrontal Cortex | 11.25_(1,66)_ | | .0013_(.293)_ | .036_(1,75)_ | .851_.568)_ | .224_(1,69)_ | .638_(.953)_ | .468⊥ |
| Ventrolateral Prefrontal Cortex | 14.93_(1,65)_ | | **<.001_(<.001)_** | 9.91_(1,79)_ | **.002_(.035)_** | 1.48_(1,65)_ | .228_(.494)_ | .313⊥ |
| Cerebellum | 5.60_(1,125)_ | | .019_(.816)_ | 1.09_(1,125)_ | .299_(.607)_ | 15.03_(1,125)_ | <.001_(.060)_ | .204∩ |
| Retrosplenial Cortex | .01_(1,125)_ | | .936_(.877)_ | 7.06_(1,125)_ | .009_(.568)_ | 14.40_(1,137)_ | <.001_(.060)_ | .150∩ |
| Precuneus | 4.51_(1,65)_ | | .038_(.790)_ | 13.41_(1,79)_ | .0005_(.330)_ | 5.60_(1,65)_ | .021_(.194)_ | .171∩ |
| Lateral Occipital Cortex | 8.81_(1,64)_ | | .004_(.095)_ | 16.04_(1,77)_ | **.0001_(.02)_** | 4.73_(1,64)_ | .033_(.060)_ | .290⊥ |

*Notes.* *Notes.* Subject was included as random effect. Group (children, young adults), Session (Day 1 remote > recent, Day 14 remote > recent), and their interaction were included as fixed effect. The following reference levels where used: for Session – Day 1; for Group – Children; F – F-value; DF – degrees of freedom; p – p-value; FDR_adj – False Discovery Rate adjusted; R2 – amount of variance explained by the model (∩- marginal; ⊥ - conditional). Type III Analysis of Variance Table with Satterthwaite’s method. *p < .05; ** < .01, *** < .001 (significant difference). All p-values of main and interactions effects were FDR-adjusted for multiple comparisons.

**Table S9.2**

*Statistical overview of the main and interaction effects of the linear mixed effects models* ***for remote > recent*** *univariate results for correctly recognized items (based on the pipeline without global signal).*

|  | **Main Effect**  **of Group** | | **Main Effect**  **of Session** | | **Group x Session Interaction** | |  |
| --- | --- | --- | --- | --- | --- | --- | --- |
| ***Regions of Interest*** | *F_(DF)_* | *p* | *F_(DF)_* | *p* | *F_(DF)_* | p | *R2* |
| Hippocampus Anterior | .002_(1,161)_ | .963_(.956)_ | .081_(1,161)_ | .776_(.862)_ | .034_(1,161)_ | .853_(.930)_ | .01∩ |
| Hippocampus Posterior | .151_(1,161)_ | .698_(.893)_ | .528_(1,161)_ | .468_(.669)_ | .006_(1,161)_ | .930_(.930)_ | .04∩ |
| Parahippocampal Gyrus Anterior | .153_(1,161)_ | .696_(.893)_ | .01_(1,161)_ | .936_(.936)_ | .013_(1,161)_ | .910_(.930)_ | .013∩ |
| Parahippocampal Gyrus Posterior | .898_(1,83)_ | .345_(.690)_ | 3.31_(1,85)_ | .071_(.236)_ | 3.03_(1,85)_ | .085_(.230)_ | .052⊥ |
| Medial Prefrontal Cortex | 2.20_(1,83)_ | .142_(.460)_ | 2.38_(1,85)_ | .125_(.213)_ | .374_(1,85)_ | .542_(.774)_ | .083⊥ |
| Ventrolateral Prefrontal Cortex | 20.37_(1,80)_ | **<.001_(<.001)_** | 8.70_(1,107)_ | **.004_(.02)_** | .921_(1,83)_ | .340_(.675)_ | .269⊥ |
| Cerebellum | 0.08_(1,161)_ | .784_(.893)_ | 1.64_(1,161)_ | .202_(.404)_ | 3.07_(1,161)_ | .082_(.230)_ | .063∩ |
| Retrosplenial Cortex | .06_(1,161)_ | .804_(.893)_ | .084_(1,161)_ | .773_(.862)_ | 2.87_(1,161)_ | .092**_(.230)_** | .036∩ |
| Precuneus | 1.78_(1,161)_ | .184_(.460)_ | .568_(1,161)_ | .452_(.669)_ | .696_(1,161)_ | .405_(.675)_ | .045∩ |
| Lateral Occipital Cortex | 4.53_(1,88)_ | .036_(.180)_ | 13.64_(1,111)_ | **<.001_(<.001)_** | 3.62_(1,88)_ | .060_(.230)_ | .297⊥ |

|  | **Main Effect**  **of Sex** | | **Main Effect**  **of Handedness** | | **Main Effect**  **of IQ** | | **Main Effect**  **Of Reaction Time** | |
| --- | --- | --- | --- | --- | --- | --- | --- | --- |
| ***Regions of Interest*** | *F_(DF)_* | *p* | *F_(DF)_* | *p* | *F_(DF)_* | p | *F_(DF)_* | *p* |
| Hippocampus Anterior | .011_(1,161)_ | .9155 | .681_(1,161)_ | .508 | .036_(1,161)_ | .850 | .293_(1,161)_ | .853 |
| Hippocampus Posterior | .039_(1,161)_ | .843 | 2.73_(1,161)_ | .068 | .002_(1,161)_ | .967 | .01_(1,161)_ | .940 |
| Parahippocampal Gyrus Anterior | .178_(1,161)_ | .674 | .828_(1,161)_ | .439 | .027_(1,161)_ | .870 | .027_(1,161)_ | .870 |
| Parahippocampal Gyrus Posterior | .042_(1,84)_ | .838 | .785_(1,93)_ | .459 | .000_1,84)_ | .987 | .289_(1,155)_ | .598 |
| Medial Prefrontal Cortex | .038_(1,87)_ | .846 | 2.93_(1,95)_ | .058 | 1.71_(1,87)_ | .194 | .090_(1,160)_ | .765 |
| Ventrolateral Prefrontal Cortex | .008_(1,81)_ | .928 | 1.04_(1,90)_ | .357 | .625_(1,81)_ | .431 | .029_(1,149)_ | .864 |
| Cerebellum | 1.22_(1,161)_ | .270 | .537_(1,161)_ | .585 | .0001_(1,161)_ | .991 | 1.07_(1,161)_ | .303 |
| Retrosplenial Cortex | .309_(1,161)_ | .579 | .749_(1,161)_ | .474 | .662_(1,161)_ | .417 | .310_(1,161)_ | .579 |
| Precuneus | .323_(1,161)_ | .570 | .856_(1,161)_ | .427 | .621_(1,161)_ | .432 | .007_(1,161)_ | .934 |
| Lateral Occipital Cortex | .001_(1,88)_ | .977 | 1.13_(1,96)_ | .327 | **7.40_(1,87)_** | **.008** | .000_(1,159)_ | .986 |

*Notes.* *Notes.* Subject was included as random effect. Group (children, young adults), Session (Day 1 remote > recent, Day 14 remote > recent), and their interaction were included as fixed effect. The following reference levels where used: for Session – Day 1; for Group – Children; F – F-value; DF – degrees of freedom; p – p-value; FDR_adj – False Discovery Rate adjusted; R2 – amount of variance explained by the model (∩- marginal; ⊥ - conditional). Type III Analysis of Variance Table with Satterthwaite’s method. *p < .05; ** <.01, ***<.001 (significant difference). All p-values of main and interactions effects were FDR-adjusted for multiple comparisons.

Table S10

*Statistical overview of the main and interaction effects of the linear mixed effects model for scene-specific reinstatement.*

|  | **Main Effect**  **of Group** | | **Main Effect**  **of Session** | | **Group x Session Interaction** | | **Main Effect**  **of BOLD activation** | |  |
| --- | --- | --- | --- | --- | --- | --- | --- | --- | --- |
| ***Regions of Interest*** | *F_(DF)_* | *p* | *F_(DF)_* | *p* | *F_(DF)_* | *p* | *F_(DF)_* | *p* | *R2* |
| HCa | 27.21_(1,86)_ | **<.001** | 100.70_(2,159)_ | **<.001** | .94_(2,159)_ | .393 | .92_(1,226)_ | .339 | .411 |
| HCp | 27.19_(1,87)_ | **<.001** | 98.18_(2,159)_ | **<.001** | 1.71(_2,158)_ | .183 | .97_(1,240)_ | .324 | .417 |
| PHGa | 23.14_(1,87)_ | **<.001** | 97.74_(2,159)_ | **<.001** | 1.62_(2,159)_ | .201 | 1.05_(1,221)_ | .307 | .397 |
| PHGp | 15.70_(1,82)_ | **<.001** | 94.40_(2,163)_ | **<.001** | 1.85(_2,155)_ | .161 | .25_(1,240)_ | .619 | .371 |
| mPFC | 8.89_(1,90)_ | **.0044** | 72.811_(2,161)_ | **<.001** | .935_(2,152)_ | .395 | 2.24_(1,221)_ | .136 | .634 |
| vlPFC | 15.18_(1,90)_ | **<.001** | 71.36_(2,172)_ | **<.001** | 1.23_(2,165)_ | .295 | .003_(1,242)_ | .955 | .591 |
| CE | 9.54_(1,87)_ | **.0038** | 59.99_(2,166)_ | **<.001** | 1.17_(2,162)_ | .313 | .679_(1,228)_ | .411 | .520 |
| RSC | 9.27_(1,89)_ | **.0038** | 79.40_(2,162)_ | **<.001** | 1.86_(2,162_ | .159 | .101_(1,242)_ | .751 | .564 |
| PC | 11.35_(1,85)_ | **.0016** | 74.33_(2,161)_ | **<.001** | 1.57_(1,160)_ | .190 | .008_(1,223)_ | .925 | .580 |
| LOC | 1.22_(1,100)_ | .271 | 64.96_(2,167)_ | **<.001** | 1.05_(2,162)_ | .350 | 1.33(_1,220)_ | .249 | .523 |

*Notes.* Subject was included as a random effect. Group (children, young adults), Delay ( recent, remote (Day 1), remote (Day 14)), and their interaction were included as fixed effect. The following reference levels where used: for Delay, recent; for Group, Children; F – F-value; DF – degrees of freedom; p – p-value; FDR_adj – False Discovery Rate adjusted; R2 – amount of variance explained by the model (Stoffel et al., 2021); mPFC – medial prefrontal cortex; vlPFC – ventrolateral prefrontal cortex; HCa – anterior hippocampus; HCp – posterior hippocampus; PHGa – anterior parahippocampal cortex; PHGp – posterior parahippocampal cortex; CE – cerebellum; PC – precuneus; RSC – retrosplenial cortex; LOC – lateral occipital cortex.. Type III Analysis of Variance Table with Satterthwaite's method. *p < .05; ** < .01, *** < .001 (significant difference). All main and interactions p-values were FDR-adjusted for multiple comparisons. All main and interactions p-values were FDR-adjusted for multiple comparisons.

Table S10.1

*Statistical overview of the main and interaction effects of the linear mixed effects model for object-specific reinstatement.*

| **Object-specific Reinstatement** | | |  |
| --- | --- | --- | --- |
| *Predictors* | *Estimate* | *CI* | *p-value* |
| (Intercept) | .47  .03  .00  -.01  .00  .00  -.01  .02 | .45 – .48 | **<0.001** |
| Group (adults vs children) |  | .01 – .05 | .013 |
| Session (Day 1 vs Day 14) |  | -.02 – .02 | .765 |
| Condition (recent vs remote) |  | -.03 – .01 | .216 |
| Group × Session |  | **-.03 – .03** | **.927** |
| Group × Condition |  | -.02 – .03 | .817 |
| Session × Condition |  | -.04 – .01 | .343 |
| Group ×Session × Condition |  | -.02 – .06 | .350 |
| **Random Effects** |  | | |
| σ^2^ | .00 | | |
| τ_00_ _subNo_ | .00 | | |
| ICC | .16 | | |
| N _subNo_ | 83 | | |
| Observations | 3058 | | |
| Marginal R^2^ / Conditional R^2^ | .401/ .498 | | |

*Notes.* CI – confidence interval; p – p-value; σ2 – residuals, τ00 – variance of the random intercept. Type III Analysis of Variance Table with Satterthwaite's method. *p < .05; ** < .01, *** < .001 (significant difference). The output table has been shortened for clarity; all interaction effects, including those involving ROI, were excluded as they did not yield significant results.

Table S10.2

*Statistical overview of the main and interaction effects of the linear mixed effects model for remote object-specific reinstatement.*

|  | **Main Effect**  **of Session** | | **Group x Session Interaction** | |
| --- | --- | --- | --- | --- |
| ***Regions of Interest*** | *F_(DF)_* | *p* | *F_(DF)_* | *p* |
| HCa | .44_(1,77)_ | .506 | .001_(1,77)_ | .973 |
| HCp | 1.09_(1,77)_ | .299 | .56(_1,77)_ | .456 |
| PHGa | .48_(1,83)_ | .487 | .946_(1,80)_ | .333 |
| PHGp | .395_(1,78)_ | .532 | .486(_1,77)_ | .488 |
| mPFC | 1.225_(1,79)_ | .272 | 1.501_(1,78)_ | .224 |
| vlPFC | 1.058_(1,82)_ | .307 | 2.011_(1,82)_ | .160 |
| CE | 424_(1,153)_ | .516 | 6.03_(1,53)_ | .015 |
| RSC | 2.106_(1,153)_ | .149 | 1.09_(1,153)_ | .289 |
| PC | 3.60_(1,153)_ | .059 | .133_(1,153)_ | .716 |
| LOC | 2.006_(1,74)_ | .161 | 1.074_(1,74)_ | .304 |

*Notes.* Subject was included as a random effect. F – F-value; DF – degrees of freedom; p – p-value; mPFC – medial prefrontal cortex; vlPFC – ventrolateral prefrontal cortex; HCa – anterior hippocampus; HCp – posterior hippocampus; PHGa – anterior parahippocampal cortex; PHGp – posterior parahippocampal cortex; CE – cerebellum; PC – precuneus; RSC – retrosplenial cortex; LOC – lateral occipital cortex. Type III Analysis of Variance Table with Satterthwaite's method. *p < .05; ** < .01, *** < .001 (significant difference).

Table S10.3

*Statistical overview of the main and interaction effects of the linear mixed effects model for scene-specific reinstatement for corpus callosum subregions.*

|  | ***Session*** | | ***Group*** | |
| --- | --- | --- | --- | --- |
| ***ROI*** | ***F_(DF)_*** | ***p*** | ***F_(DF)_*** | ***p*** |
| Corpus Collosum – corpus | 3.88_(2,367)_ | **.021** | 1.33_(1,367)_ | .248 |
| Corpus Collosum – gernu | 3.87_(2,373)_ | **.022** | 1.69_(1,373)_ | .195 |
| Corpus Collosum – splenium | 3.88_(2,373)_ | **.022** | .405_(1,373)_ | .525 |

*Notes.* Subject was included as a random effect. Group (children, young adults), Session ( recent, remote (Day 1), remote (Day 14)) as fixed effect. The following reference levels where used: for Session, recent; for Group, Children; F – F-value; DF – degrees of freedom; p – p-value; Type III Analysis of Variance Table with Satterthwaite's method. *p < .05; ** < .01, *** < .001 (significant difference).

Table S10.4

*Statistical overview of LME-model based Sidak corrected post hoc comparisons for scene-specific reinstatement analysis for corpus callosum subregions (based on LME-model described in Table S10.3).*

|  | Recent > Remote Day1 | | | Remote Day 1 > Day 14 | | |
| --- | --- | --- | --- | --- | --- | --- |
|  | *b* | *t_(DF)_* | *p* | *b* | *t_(DF)_* | *p* |
| Corpus Collosum-Corpus | .029 | 2.23_(156)_ | .080 | .024 | 1.78_(163)_ | .213 |
| Corpus Collosum - gernu | .027 | 2.05_(159)_ | .121 | .028 | 2.04_(165)_ | .124 |
| Corpus Collosum- splenium | .026 | 2.03_(159)_ | .128 | .026 | 1.94_(166)_ | .147 |

*Notes.* Degrees of freedom were adjusted based on Kenward-Roger methods. b – Beta values; t – t-value; DF – degrees of freedom; p – p-value; *p < .05; ** < .01, *** < .001 (significant difference).

Figure S4

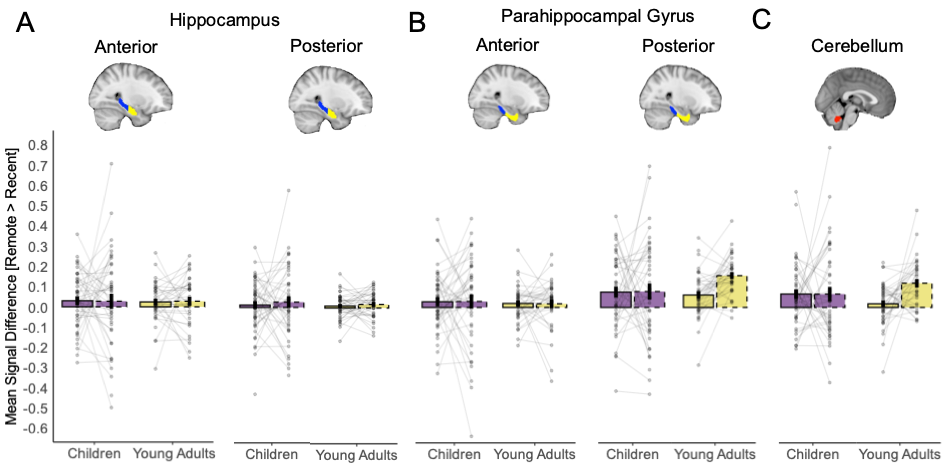

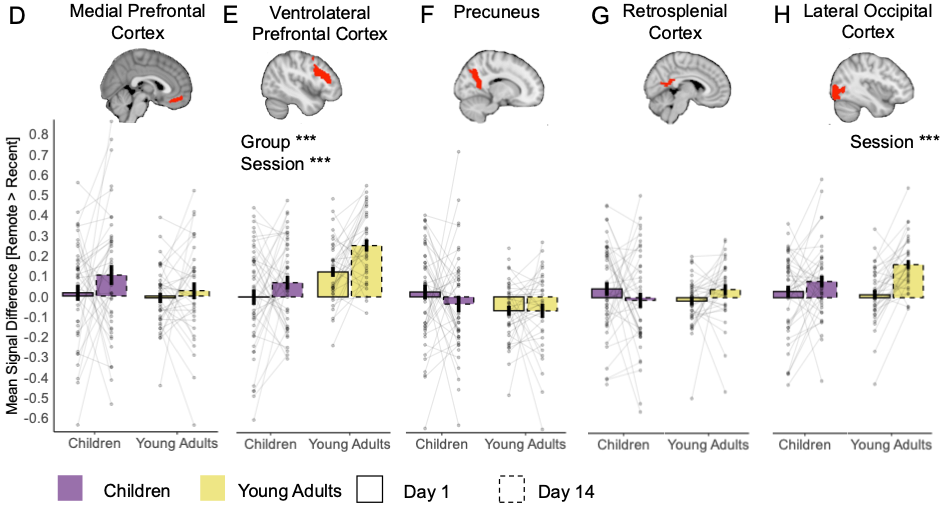

**Mean Signal Differences Between Correct Remote and Recent Memories (without global signal in preprocessing).** The figure presents mean signal difference for remote > recent contrast across sessions and groups during the object presentation time window in **(A)** Anterior and Posterior Hippocampus; (**B**) Anterior and Posterior Parahippocampal Gyrus; **(C)** Cerebellum; **(D)** Medial Prefrontal Cortex; **(E)** Ventrolateral Prefrontal Cortex; **(F)** Precuneus; **(G)** Retrosplenial Cortex; **(H)** Lateral Occipital Cortex. *Note:* Bars indicate the group mean for each session (solid lines for Day 1, dashed lines for Day 14), plotted separately for children and young adults. Error bars represent ±1 standard error of the mean. The colour indicated the age groups: purple for children and khaki yellow for young adults. Across all panels, mean of individual subject data are shown with transparent points. The connecting faint lines reflect within-subject differences across sessions. Orange asterisks denote significant difference of **remote > recent** contrast from zero. An upward orange arrow indicates that this difference is greater than zero, while a downward arrow indicates that this is less than zero. **p* < .05; ***p* < .01; ****p* < .001(significant difference); non-significant differences were not specifically highlighted. Significant main and interaction effects are highlighted by the corresponding asterisks. All main and interaction p-values were FDR-adjusted for multiple comparisons.

Figure S5

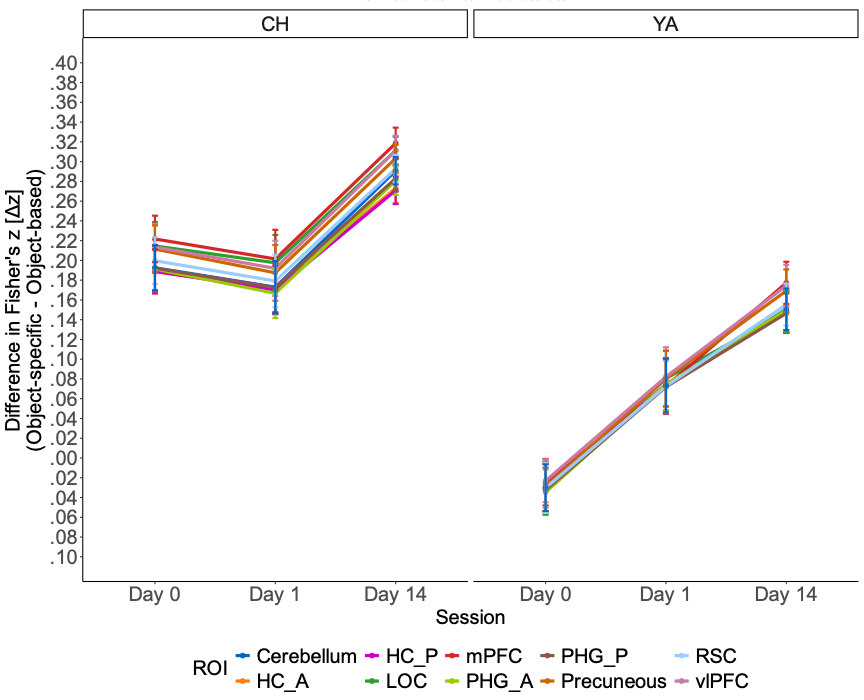

**Object-specific neural reinstatement**

Object-specific neural reinstatement defined as the difference between Fisher-transformed scene-specific and set-specific representational similarity (for incorrectly remembered items). Scene-specific neural reinstatement index by group (children vs. adults) and session (Day 0 – recent, Day 1 – remote short delay, Day 14 – remote long delay). HC_A – hippocampus anterior; HC_P – hippocampus posterior; PHG_A – parahippocampal gyrus anterior; PHG_P – parahippocampal gyrus posterior; mPFC – medial prefrontal cortex; vlPFC – ventrolateral prefrontal cortex; RSC – retrosplenial cortex; LOC – lateral occipital cortex. Error bars indicate standard error.

Figure S6

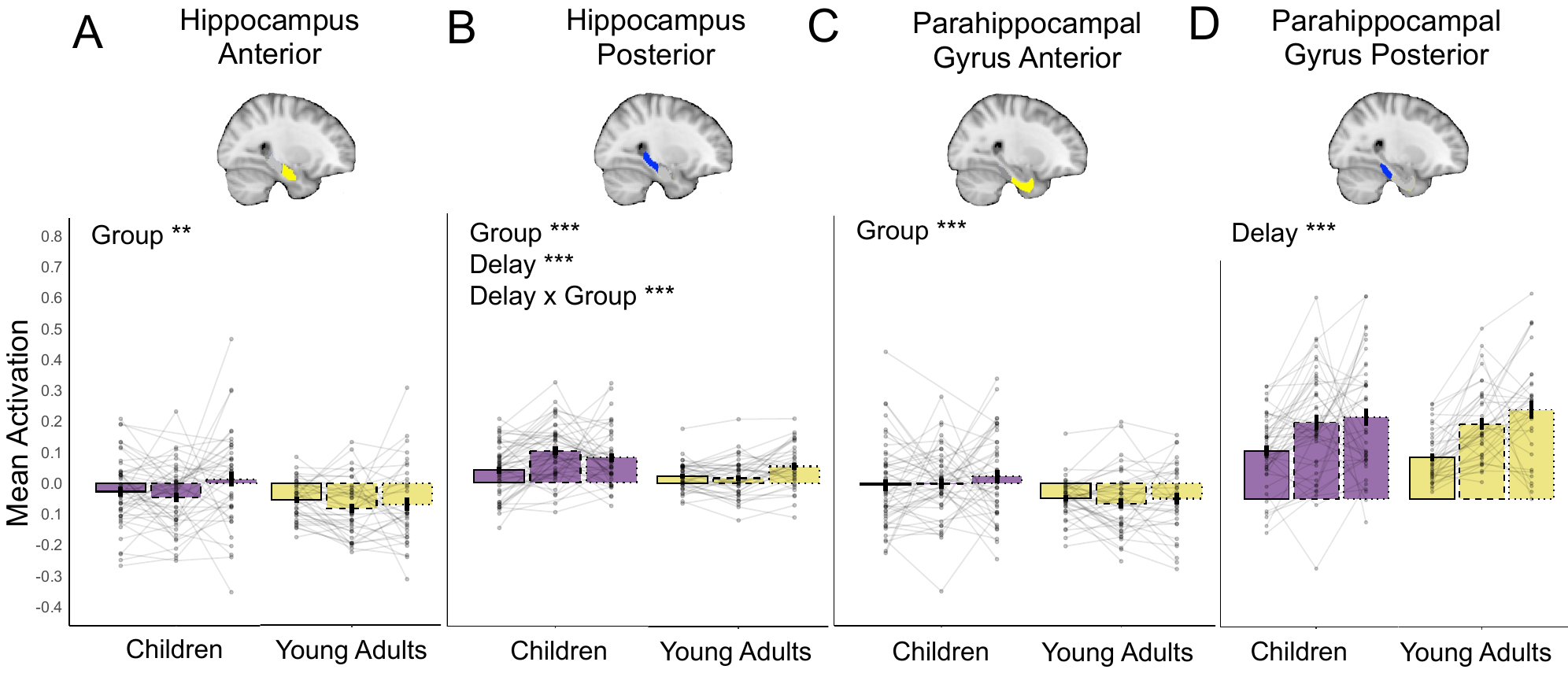

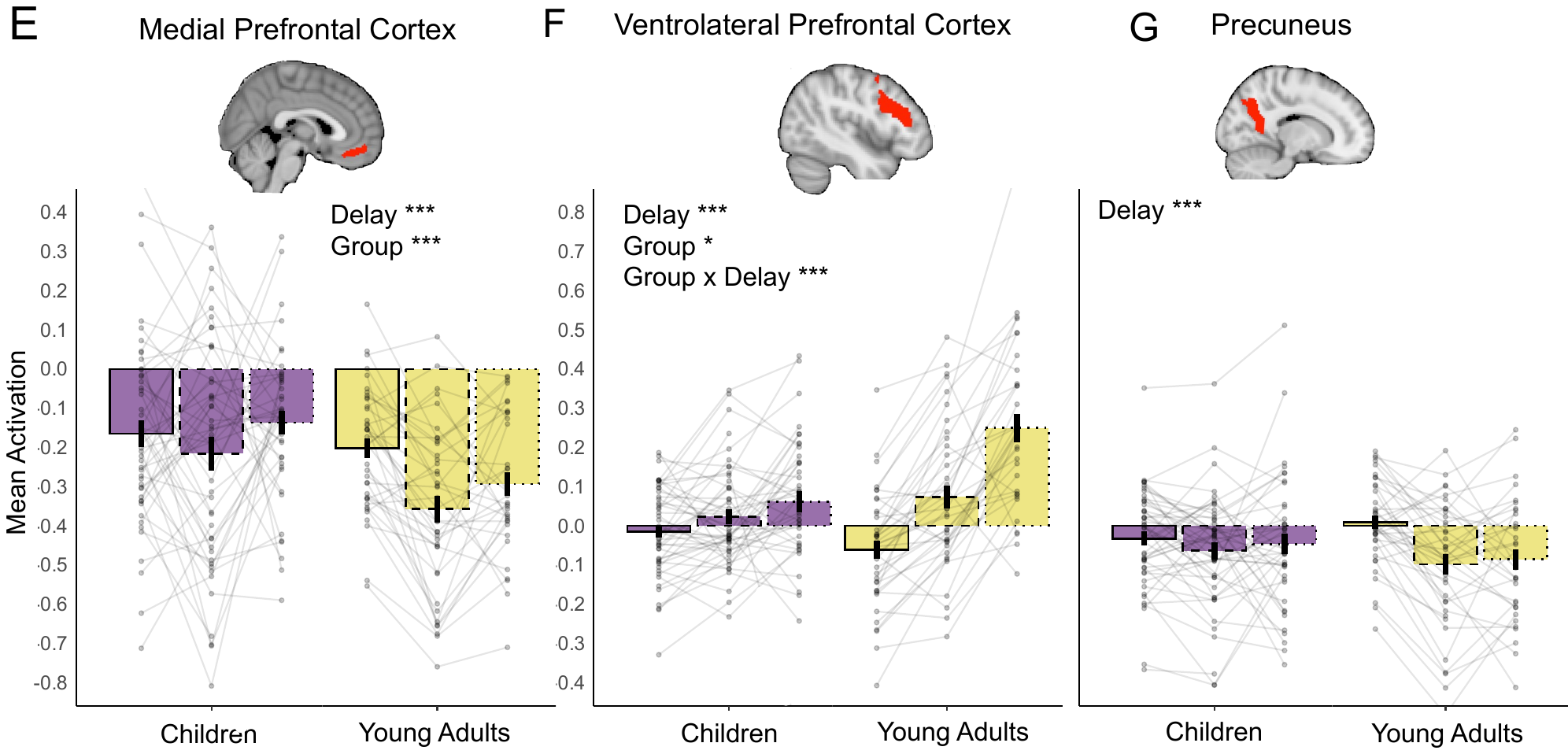

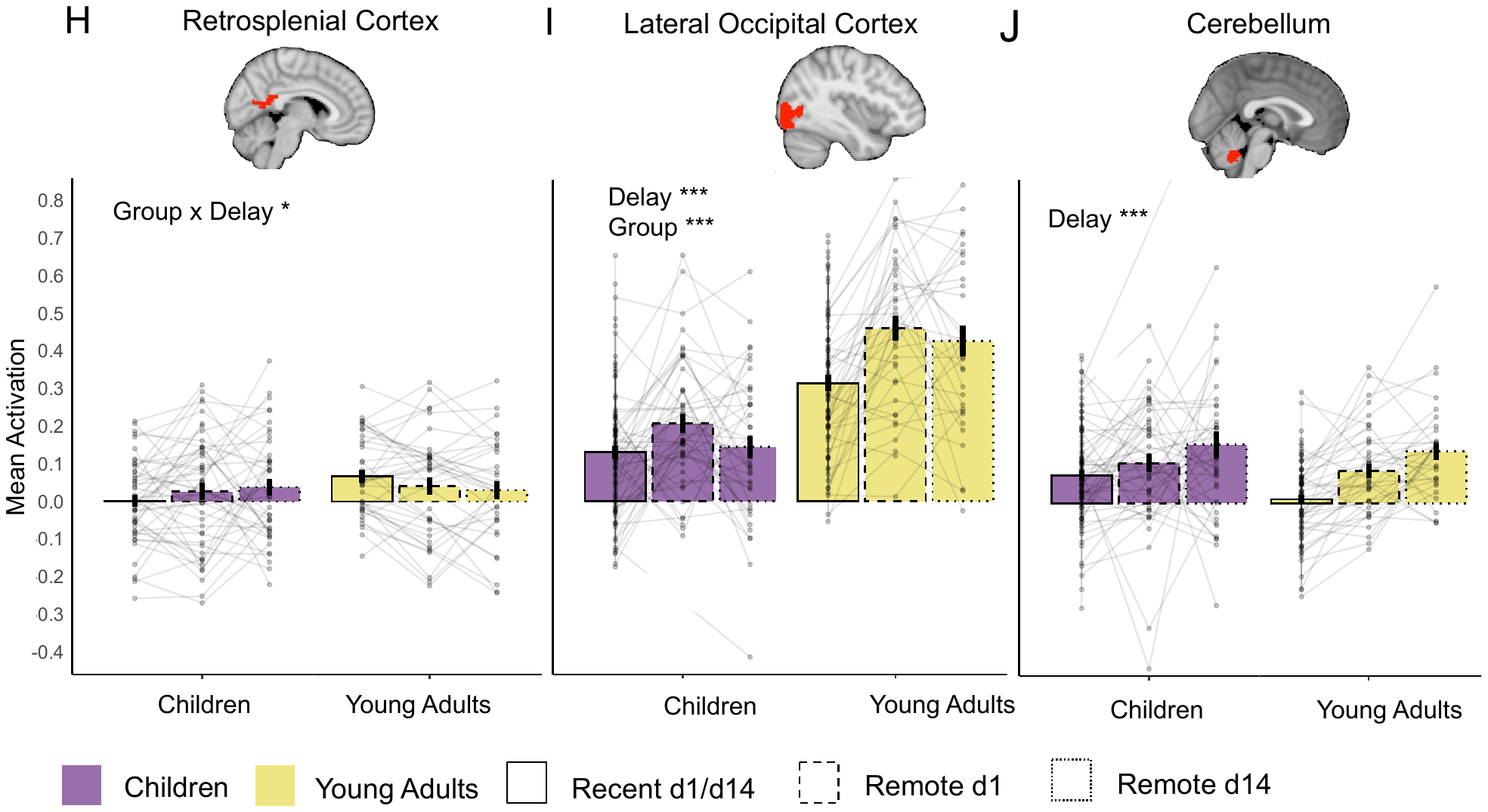

**Mean Neural Activation for Correctly Recalled Memories during Scene Presentation Time Window.**

The figure presents mean signal intensity for correctly recalled recent, short delay remote and long delay remote memories in children and adults in **(A)** anterior hippocampus; **(B)** posterior hippocampus; (**C**) anterior parahippocamla gyrus; **(D)** posterior parahippocampal gyrus; **(E)** medial prefrontal cortex; **(F)** ventrolateral prefrontal cortex; **(G)** precuneus; **(H)** retrosplenial cortex; **(I)** lateral occipital cortex; **(J)** cerebellum. *Note:* Bars represent the average signal difference. The colour indicated the age groups: purple for children and khaki yellow for young adults. Solid-lined bars represent data from Day 1, while dashed-lined bars depict data from Day 14. Across all panels, mean of individual subject data are shown with transparent points. The connecting faint lines reflect within-subject differences across sessions. Error bars indicate standard error of the mean. **p* < .05; ***p* < .01; ****p* < .001(significant difference); non-significant differences were not specifically highlighted. Significance main and interaction effects are highlighted by the corresponding asterisks. All main and interactions p-values were FDR-adjusted for multiple comparisons.

Table S11

*Statistical overview of the main and interaction effects of the linear mixed effects model for scene-based univariate neural analysis*

|  | **Main Effect**  **of Group** | | **Main Effect**  **of Delay** | | **Group x Delay Interaction** | |  |
| --- | --- | --- | --- | --- | --- | --- | --- |
| ***Regions of Interest*** | *F_(DF)_* | p | *F_(DF)_* | *p* | *F_(DF)_* | *p* | *R2* |
| HCa | **7.16(1,94)** | **.009** | 2.35(2,238) | .097 | 3.02(2,238) | .051 | .320 |
| HCp | **11.67(1,97)** | **.0009** | **8.25(2,241)** | **.0003** | **7.19(2,241)** | **.0009** | .374 |
| PHGa | **11.02(1,90)** | **.001** | .42(2,234) | .660 | .927(2,234) | .397 | .326 |
| PHGp | .012(1,95) | .914 | **36.46(2,240)** | **<.001** | .749(2,240) | .474 | .377 |
| Medial Prefrontal Cortex | **10.28_(1,85)_** | **.002** | **7.21_(2,163)_** | **<.001** | 2.94 _(2,163)_ | .056 | .105 |
| Ventrolateral Prefrontal Cortex | **5.96_(1,88)_** | **.016** | **55.14_(2,164)_** | **<.001** | **20.47_(2,164)_** | **<.001** | .262 |
| Cerebellum | 1.98_(1,80)_ | .163 | **13.63_(2,158)_** | **<.001** | .065_(2,158)_ | .522 | .084 |
| Retrosplenial Cortex | 1.05_(1,88)_ | .308 | .00_(2,164)_ | .999 | **3.28_(2,164)_** | **.039** | .023 |
| Precuneus | .19_(1,88)_ | .666 | **12.01_(2,163)_** | **<.001** | **4.54_(2,163)_** | **.012** | .056 |
| Lateral Occipital Cortex | **54.52_(1,88)_** | **<.001** | **17.09_(2,163)_** | **<.001** | **3.55_(2,163)_** | **.031** | .338 |

*Notes.* Subject was included as random effect. Group (children, young adults), Delay ( recent, remo te (Day 1), remote (Day 14)), and their interaction were included as fixed effect. The following reference levels where used: for Delay, recent; for Group, Children; mPFC – medial prefrontal cortex; vlPFC – ventrolateral prefrontal cortex; HCa – anterior hippocampus; HCp – posterior hippocampus; PHGa – anterior parahippocampal cortex; PHGp – posterior parahippocampal cortex;CE – cerebellum; PC – precuneus; RSC – retrosplenial cortex; LOC – lateral occipital cortex. F – F-value; DF – degrees of freedom; p – p-value; R2 – amount of variance explained by the model (Stoffel et al., 2021). Type III Analysis of Variance Table with Satterthwaite's method. *p < .05; ** < .01, *** < .001 (significant difference).

Table S12

*Test of gist-like representations index for significance (higher than zero).*

|  |  | Recent Pre-activation | | | Short-Delay Pre-activation | | | | | | Long-Delay Pre-activation | | | | | | | |
| --- | --- | --- | --- | --- | --- | --- | --- | --- | --- | --- | --- | --- | --- | --- | --- | --- | --- | --- |
|  |  | **Children** | | | | | | | | | | | | | | | | |
| **ROI** | *mean* | | *p* | *p_(FDRadj)_* | | *mean* | *p* | | | *p_(FDRadj)_* | | | | *mean* | | *p* | | *p _(FDRadj)_* |
| mPFC | .006 | | .135 | .303 | | .003 | .282 | | | .346 | | | **.026** | | | **.002** | | **.013** |
| vlPFC | -.001 | | .583 | .583 | | .006 | .088 | | | .178 | | | **.020** | | | **.001** | | **.007** |
| HCa | .003 | | .045 | .135 | | .004 | .038 | | | .135 | | | -.001 | | | .586 | | .704 |
| HCp | .002 | | .076 | .196 | | .003 | .097 | | | .196 | | | .004 | | | .227 | | .340 |
| PHGa | .002 | | .225 | .415 | | .005 | .018 | | | .113 | | | .002 | | | .328 | | .415 |
| PHGp | .003 | | .127 | .254 | | .004 | .045 | | | .181 | | | -.001 | | | .617 | | .741 |
| CE | .001 | | .373 | .590 | | -.001 | .553 | | | .590 | | | .012 | | | .125 | | .590 |
| PC | .004 | | .066 | .099 | | **.009** | **.007** | | | **.044** | | | .012 | | | .042 | | .085 |
| RSC | .004 | | .019 | .090 | | .006 | .030 | | | .090 | | | .010 | | | .053 | | .105 |
| LOC | .002 | | .253 | .304 | | **.011** | **.008** | | | **.024** | | | .009 | | | .107 | | .161 |
|  |  | **Young Adults** | | | | | | | | | | | | | | | | |
|  | *mean* | | *p* | *p_(FDRadj)_* | *mean* | | | *p* | *p_(FDRadj)_* | | | *mean* | | | *p* | | *p_(FDRadj_* | |
| mPFC | .002 | | .151 | .303 | -.00002 | | | .553 | .553 | | | .002 | | | .288 | | .346 | |
| vlPFC | .001 | | .357 | .532 | .005 | | | .063 | .178 | | | .001 | | | .443 | | .532 | |
| HCa | .002 | | .116 | .232 | .001 | | | .208 | .313 | | | -.002 | | | .895 | | .900 | |
| HCp | .0003 | | .379 | .456 | .003 | | | .016 | .096 | | | -.003 | | | .882 | | .883 | |
| PHGa | .002 | | .082 | .245 | .001 | | | .346 | .415 | | | -.004 | | | .909 | | .909 | |
| PHGp | .001 | | .189 | .285 | .003 | | | .060 | .181 | | | -.004 | | | .939 | | .940 | |
| CE | .0002 | | .455 | .590 | .001 | | | .278 | .590 | | | -.0008 | | | .590 | | .590 | |
| PC | .002 | | .125 | .150 | .004 | | | .032 | .085 | | | -.001 | | | .567 | | .568 | |
| RSC | .002 | | .070 | .106 | .002 | | | .097 | .116 | | | -.001 | | | .678 | | .679 | |
| LOC | .004 | | .039 | .078 | **.006** | | | **.004** | **.024** | | | -.002 | | | .779 | | .779 | |

*Notes*.To test for significance we used one-sample permutation t-test for more robust calculations with Monte-Carlo permutation percentile confidence interval. The p-values of child group were corrected for False Discovery Rate (FDR) for multiple comparisons. ROI – region of interest; p – p-value; FDRadj – False Discovery Rate adjustment; mPFC – medial prefrontal cortex; vlPFC – ventrolateral prefrontal cortex; HCa – anterior hippocampus; HCp – posterior hippocampus; PHGa – anterior parahippocampal cortex; PHGp – posterior parahippocampal cortex; CE – cerebellum; PC – precuneus; RSC – retrosplenial cortex; LOC – lateral occipital cortex. *p < .05; ** < .01, *** < .001 (significant difference).

Table S12.1

*Test of gist-like reinstatement index for significance (higher than zero) based on cross-run comparisons.*

|  |  | Recent Pre-activation | | | Short-Delay Pre-activation | | | | | | Long-Delay Pre-activation | | | | | | | |
| --- | --- | --- | --- | --- | --- | --- | --- | --- | --- | --- | --- | --- | --- | --- | --- | --- | --- | --- |
|  |  | **Children** | | | | | | | | | | | | | | | | |
| **ROI** | *mean* | | *p* | *p_(FDRadj)_* | | *mean* | *p* | | | *p_(FDRadj)_* | | | | *mean* | | *p* | | *p _(FDRadj)_* |
| mPFC | .001 | | .416 | .807 | | -.011 | .988 | | | **.**988 | | | .014 | | | **.037** | | .126 |
| vlPFC | -.008 | | .897 | .897 | | .002 | .327 | | | .491 | | | .022 | | | **.0007** | | **.004** |
| HCa | .002 | | .319 | .891 | | -.000 | .542 | | | .891 | | | -.005 | | | .891 | | .891 |
| HCp | .002 | | .112 | .335 | | -.001 | .569 | | | .853 | | | .004 | | | .167 | | .334 |
| PHGa | -.003 | | .834 | .942 | | .004 | .117 | | | .700 | | | -.003 | | | .714 | | .942 |
| PHGp | -.001 | | .570 | .855 | | .000 | .477 | | | .855 | | | -.004 | | | .781 | | .862 |
| CE | -.002 | | .770 | .913 | | -.004 | .721 | | | .913 | | | .014 | | | .090 | | .539 |
| PC | .001 | | .312 | .375 | | .005 | .187 | | | .281 | | | .012 | | | .053 | | .135 |
| RSC | -.000 | | .576 | .691 | | .003 | .247 | | | .383 | | | .004 | | | .255 | | .383 |
| LOC | .003 | | .290 | .397 | | .012 | **.022** | | | .066 | | | .004 | | | .330 | | .396 |
|  |  | **Young Adults** | | | | | | | | | | | | | | | | |
|  | *mean* | | *p* | *p_(FDRadj)_* | *mean* | | | *p* | *p_(FDRadj)_* | | | *mean* | | | *p* | | *p_(FDRadj_* | |
| mPFC | .004 | | **.042** | .127 | -.001 | | | .568 | .807 | | | -.002 | | | .672 | | .807 | |
| vlPFC | .003 | | .161 | .322 | .009 | | | .**032** | .097 | | | .001 | | | .432 | | .518 | |
| HCa | -.001 | | .307 | .891 | -.001 | | | .759 | .891 | | | -.001 | | | .758 | | .891 | |
| HCp | -.001 | | .842 | .884 | -.001 | | | .052 | .309 | | | -.002 | | | .884 | | .884 | |
| PHGa | -.000 | | .528 | .942 | .001 | | | .398 | .942 | | | -.001 | | | .923 | | .942 | |
| PHGp | .001 | | .229 | .697 | .003 | | | .067 | .395 | | | -.002 | | | .862 | | .862 | |
| CE | -.003 | | .912 | .913 | -.002 | | | .796 | .913 | | | -.001 | | | .332 | | .913 | |
| PC | .004 | | **.028** | .135 | .004 | | | .067 | .135 | | | -.002 | | | .699 | | .699 | |
| RSC | .003 | | **.011** | .070 | .002 | | | .082 | .247 | | | -.001 | | | .773 | | .723 | |
| LOC | .002 | | .122 | .245 | **.007** | | | **.007** | **.042** | | | -.007 | | | .857 | | .857 | |

*Notes*.To test for significance we used one-sample permutation t-test for more robust calculations with Monte-Carlo permutation percentile confidence interval. ROI – region of interest; p – p-value; FDRadj – False Discovery Rate adjustment; mPFC – medial prefrontal cortex; vlPFC – ventrolateral prefrontal cortex; HCa – anterior hippocampus; HCp – posterior hippocampus; PHGa – anterior parahippocampal cortex; PHGp – posterior parahippocampal cortex; CE – cerebellum; PC – precuneus; RSC – retrosplenial cortex; LOC – lateral occipital cortex. *p < .05; ** < .01, *** < .001 (significant difference).

Table S12.2

*Test of gist-like reinstatement index for significance (higher than zero) for within run comparisons.*

|  |  | Recent Pre-activation | | | Short-Delay Pre-activation | | | | | | Long-Delay Pre-activation | | | | | | | |
| --- | --- | --- | --- | --- | --- | --- | --- | --- | --- | --- | --- | --- | --- | --- | --- | --- | --- | --- |
|  |  | **Children** | | | | | | | | | | | | | | | | |
| **ROI** | *mean* | | *p* | *p_(FDRadj)_* | | *mean* | *p* | | | *p_(FDRadj)_* | | | | *mean* | | *p* | | *p _(FDRadj)_* |
| mPFC | .010 | | .084 | .168 | | **.028** | **.008** | | | **.034** | | | **.029** | | | **.011** | | **.034** |
| vlPFC | .007 | | .068 | .135 | | .010 | .038 | | | **.**135 | | | .016 | | | .047 | | .135 |
| HCa | .006 | | .035 | .104 | | **.009** | **.003** | | | **.018** | | | .003 | | | .320 | | .385 |
| HCp | .002 | | .203 | .311 | | .007 | .013 | | | .079 | | | .002 | | | .381 | | .458 |
| PHGa | .006 | | .035 | .105 | | .008 | .012 | | | .069 | | | .007 | | | .196 | | .294 |
| PHGp | .006 | | .062 | .186 | | .008 | .023 | | | .140 | | | -.001 | | | .560 | | .672 |
| CE | .005 | | .140 | .280 | | .005 | .191 | | | .287 | | | -.0004 | | | .516 | | .620 |
| PC | .007 | | .059 | .179 | | **.012** | **.007** | | | **.042** | | | .006 | | | .223 | | .334 |
| RSC | .008 | | .008 | .050 | | .009 | .025 | | | **.**076 | | | .010 | | | .083 | | .166 |
| LOC | .002 | | .305 | .365 | | .010 | .045 | | | .140 | | | .009 | | | .164 | | .246 |
|  |  | **Young Adults** | | | | | | | | | | | | | | | | |
|  | *mean* | | *p* | *p_(FDRadj)_* | *mean* | | | *p* | *p_(FDRadj)_* | | | *mean* | | | *p* | | *p_(FDRadj_* | |
| mPFC | .0003 | | .450 | .502 | -.00002 | | | .502 | .502 | | | .006 | | | .126 | | .189 | |
| vlPFC | -.003 | | .786 | .787 | -.0003 | | | .535 | .712 | | | .001 | | | .593 | | .712 | |
| HCa | .003 | | .115 | .172 | .004 | | | .094 | .172 | | | -.004 | | | .863 | | .863 | |
| HCp | .002 | | .207 | .311 | .005 | | | .048 | .144 | | | -.003 | | | .822 | | .822 | |
| PHGa | .004 | | .063 | .127 | .001 | | | .369 | .443 | | | -.004 | | | .774 | | .774 | |
| PHGp | .003 | | .152 | .304 | -.005 | | | .908 | .908 | | | .006 | | | .242 | | .365 | |
| CE | .003 | | .098 | .280 | .005 | | | .063 | .280 | | | -.003 | | | .735 | | .736 | |
| PC | .0004 | | .446 | .445 | .003 | | | .124 | .247 | | | .001 | | | .382 | | .445 | |
| RSC | .003 | | .302 | .361 | .002 | | | .236 | .354 | | | -.0008 | | | .582 | | .583 | |
| LOC | .004 | | .065 | .140 | .004 | | | .070 | .140 | | | .001 | | | .403 | | .402 | |

*Notes*.To test for significance we used one-sample permutation t-test for more robust calculations with Monte-Carlo permutation percentile confidence interval. The p-values of child group were corrected for False Discovery Rate (FDR) for multiple comparisons. ROI – region of interest; p – p-value; FDRadj – False Discovery Rate adjustment; mPFC – medial prefrontal cortex; vlPFC – ventrolateral prefrontal cortex; HCa – anterior hippocampus; HCp – posterior hippocampus; PHGa – anterior parahippocampal cortex; PHGp – posterior parahippocampal cortex; CE – cerebellum; PC – precuneus; RSC – retrosplenial cortex; LOC – lateral occipital cortex. *p < .05; ** <.01, ***<.001 (significant difference).

### References

Abraham, A., Pedregosa, F., Eickenberg, M., Gervais, P., Mueller, A., Kossaifi, J., Gramfort, A., Thirion, B., & Varoquaux, G. (2014). Machine learning for neuroimaging with scikit-learn. *Frontiers in Neuroinformatics*, *8*. https://doi.org/10.3389/fninf.2014.00014

Andersson, J. L. R., Skare, S., & Ashburner, J. (2003). How to correct susceptibility distortions in spin-echo echo-planar images: application to diffusion tensor imaging. *NeuroImage*, *20*(2), 870–888. https://doi.org/10.1016/S1053-8119(03)00336-7

AVANTS, B., EPSTEIN, C., GROSSMAN, M., & GEE, J. (2008). Symmetric diffeomorphic image registration with cross-correlation: Evaluating automated labeling of elderly and neurodegenerative brain. *Medical Image Analysis*, *12*(1), 26–41. https://doi.org/10.1016/j.media.2007.06.004

Bates, D., Mächler, M., Bolker, B. M., & Walker, S. C. (2015). Fitting Linear Mixed-Effects Models Using lme4. *Journal of Statistical Software*, *67*(1), 1–48. https://doi.org/10.18637/JSS.V067.I01

Behzadi, Y., Restom, K., Liau, J., & Liu, T. T. (2007). A component based noise correction method (CompCor) for BOLD and perfusion based fMRI. *NeuroImage*, *37*(1), 90–101. https://doi.org/10.1016/j.neuroimage.2007.04.042

Cox, R. W., & Hyde, J. S. (1997). Software tools for analysis and visualization of fMRI data. *NMR in Biomedicine*, *10*(4–5), 171–178. https://doi.org/10.1002/(SICI)1099-1492(199706/08)10:4/5<171::AID-NBM453>3.0.CO;2-L

Criss, A. H. (2010). Differentiation and response bias in episodic memory: Evidence from reaction time distributions. *Journal of Experimental Psychology: Learning, Memory, and Cognition*, *36*(2), 484–499. https://doi.org/10.1037/a0018435

Esteban, O., Blair, R., Markiewicz, C. J., Berleant, S. L., Moodie, C., Ma, F., & Isik, A. I. (2018). *fMRIPrep 22.0.0.*

Esteban, O., Markiewicz, C. J., Blair, R. W., Moodie, C. A., Isik, A. I., Erramuzpe, A., Kent, J. D., Goncalves, M., DuPre, E., Snyder, M., Oya, H., Ghosh, S. S., Wright, J., Durnez, J., Poldrack, R. A., & Gorgolewski, K. J. (2019). fMRIPrep: a robust preprocessing pipeline for functional MRI. *Nature Methods*, *16*(1), 111–116. https://doi.org/10.1038/s41592-018-0235-4

Evans, A. C., Janke, A. L., Collins, D. L., & Baillet, S. (2012). Brain templates and atlases. *NeuroImage*, *62*(2), 911–922. https://doi.org/10.1016/j.neuroimage.2012.01.024

Fonov, V., Evans, A., McKinstry, R., Almli, C., & Collins, D. (2009). Unbiased nonlinear average age-appropriate brain templates from birth to adulthood. *NeuroImage*, *47*, S102. https://doi.org/10.1016/S1053-8119(09)70884-5

Forstmann, B. U., Ratcliff, R., & Wagenmakers, E.-J. (2016). Sequential Sampling Models in Cognitive Neuroscience: Advantages, Applications, and Extensions. *Annual Review of Psychology*, *67*(1), 641–666. https://doi.org/10.1146/annurev-psych-122414-033645

Fudenberg, D., Newey, W., Strack, P., & Strzalecki, T. (2020). Testing the drift-diffusion model. *Proceedings of the National Academy of Sciences*, *117*(52), 33141–33148. https://doi.org/10.1073/pnas.2011446117

Gorgolewski, K., Burns, C. D., Madison, C., Clark, D., Halchenko, Y. O., Waskom, M. L., & Ghosh, S. S. (2011). Nipype: A Flexible, Lightweight and Extensible Neuroimaging Data Processing Framework in Python. *Frontiers in Neuroinformatics*, *5*. https://doi.org/10.3389/fninf.2011.00013

Gorgolewski, K. J., Auer, T., Calhoun, V. D., Craddock, R. C., Das, S., Duff, E. P., Flandin, G., Ghosh, S. S., Glatard, T., Halchenko, Y. O., Handwerker, D. A., Hanke, M., Keator, D., Li, X., Michael, Z., Maumet, C., Nichols, B. N., Nichols, T. E., Pellman, J., … Poldrack, R. A. (2016). The brain imaging data structure, a format for organizing and describing outputs of neuroimaging experiments. *Scientific Data*, *3*(1), 160044. https://doi.org/10.1038/sdata.2016.44

Greve, D. N., & Fischl, B. (2009). Accurate and robust brain image alignment using boundary-based registration. *NeuroImage*, *48*(1), 63–72. https://doi.org/10.1016/j.neuroimage.2009.06.060

Jenkinson, M., Bannister, P., Brady, M., & Smith, S. (2002). Improved Optimization for the Robust and Accurate Linear Registration and Motion Correction of Brain Images. *NeuroImage*, *17*(2), 825–841. https://doi.org/10.1006/nimg.2002.1132

Jenkinson, M., & Smith, S. (2001). A global optimisation method for robust affine registration of brain images. *Medical Image Analysis*, *5*(2), 143–156. https://doi.org/10.1016/S1361-8415(01)00036-6

Kuznetsova, A., Brockhoff, P. B., & Christensen, R. H. B. (2017). lmerTest Package: Tests in Linear Mixed Effects Models. *Journal of Statistical Software*, *82*(13), 1–26. https://doi.org/10.18637/JSS.V082.I13

Lanczos, C. (1964). Evaluation of Noisy Data. *Journal of the Society for Industrial and Applied Mathematics Series B Numerical Analysis*, *1*(1), 76–85. https://doi.org/10.1137/0701007

Lerche, V., & Voss, A. (2019). Experimental validation of the diffusion model based on a slow response time paradigm. *Psychological Research*, *83*(6), 1194–1209. https://doi.org/10.1007/s00426-017-0945-8

Palada, H., Neal, A., Vuckovic, A., Martin, R., Samuels, K., & Heathcote, A. (2016). Evidence accumulation in a complex task: Making choices about concurrent multiattribute stimuli under time pressure. *Journal of Experimental Psychology: Applied*, *22*(1), 1–23. https://doi.org/10.1037/xap0000074

Patriat, R., Reynolds, R. C., & Birn, R. M. (2017). An improved model of motion-related signal changes in fMRI. *NeuroImage*, *144*, 74–82. https://doi.org/10.1016/j.neuroimage.2016.08.051

Power, J. D., Mitra, A., Laumann, T. O., Snyder, A. Z., Schlaggar, B. L., & Petersen, S. E. (2014). Methods to detect, characterize, and remove motion artifact in resting state fMRI. *NeuroImage*, *84*, 320–341. https://doi.org/10.1016/j.neuroimage.2013.08.048

Pruim, R. H. R., Mennes, M., van Rooij, D., Llera, A., Buitelaar, J. K., & Beckmann, C. F. (2015). ICA-AROMA: A robust ICA-based strategy for removing motion artifacts from fMRI data. *NeuroImage*, *112*, 267–277. https://doi.org/10.1016/j.neuroimage.2015.02.064

Ratcliff, R., Love, J., Thompson, C. A., & Opfer, J. E. (2012). Children Are Not Like Older Adults: A Diffusion Model Analysis of Developmental Changes in Speeded Responses. *Child Development*, *83*(1), 367–381. https://doi.org/10.1111/j.1467-8624.2011.01683.x

Ratcliff, R., & McKoon, G. (2008). The Diffusion Decision Model: Theory and Data for Two-Choice Decision Tasks. *Neural Computation*, *20*(4), 873–922. https://doi.org/10.1162/neco.2008.12-06-420

Ratcliff, R., Thapar, A., & McKoon, G. (2011). Effects of aging and IQ on item and associative memory. *Journal of Experimental Psychology: General*, *140*(3), 464–487. https://doi.org/10.1037/a0023810

Reuter, M., Rosas, H. D., & Fischl, B. (2010). Highly accurate inverse consistent registration: A robust approach. *NeuroImage*, *53*(4), 1181–1196. https://doi.org/10.1016/j.neuroimage.2010.07.020

Satterthwaite, T. D., Elliott, M. A., Gerraty, R. T., Ruparel, K., Loughead, J., Calkins, M. E., Eickhoff, S. B., Hakonarson, H., Gur, R. C., Gur, R. E., & Wolf, D. H. (2013). An improved framework for confound regression and filtering for control of motion artifact in the preprocessing of resting-state functional connectivity data. *NeuroImage*, *64*, 240–256. https://doi.org/10.1016/j.neuroimage.2012.08.052

Stoffel, M. A., Nakagawa, S., & Schielzeth, H. (2021). partR2 : partitioning R ^2^ in generalized linear mixed models. *PeerJ*, *9*, e11414. https://doi.org/10.7717/peerj.11414

Turker, H. B., & Swallow, K. M. (2022). Diffusion Decision Modeling of Retrieval Following the Temporal Selection of Behaviorally Relevant Moments. *Computational Brain & Behavior*, *5*(3), 302–325. https://doi.org/10.1007/s42113-022-00148-z

Tustison, N. J., Avants, B. B., Cook, P. A., Yuanjie Zheng, Egan, A., Yushkevich, P. A., Gee, J. C., Zheng, Y., Egan, A., Yushkevich, P. A., Gee, J. C., Yuanjie Zheng, Egan, A., Yushkevich, P. A., & Gee, J. C. (2010). N4ITK: Improved N3 Bias Correction. *IEEE Transactions on Medical Imaging*, *29*(6), 1310–1320. https://doi.org/10.1109/TMI.2010.2046908

Wagenmakers, E.-J., Van Der Maas, H. L. J., & Grasman, R. P. P. P. (2007a). An EZ-diffusion model for response time and accuracy. *Psychonomic Bulletin & Review*, *14*(1), 3–22. https://doi.org/10.3758/BF03194023

Wagenmakers, E.-J., Van Der Maas, H. L. J., & Grasman, R. P. P. P. (2007b). An EZ-diffusion model for response time and accuracy. *Psychonomic Bulletin & Review*, *14*(1), 3–22. https://doi.org/10.3758/BF03194023

Zhang, Y., Brady, M., & Smith, S. (2001). Segmentation of brain MR images through a hidden Markov random field model and the expectation-maximization algorithm. *IEEE Transactions on Medical Imaging*, *20*(1), 45–57. https://doi.org/10.1109/42.906424

Zhou, J., Osth, A. F., Lilburn, S. D., & Smith, P. L. (2021). A circular diffusion model of continuous-outcome source memory retrieval: Contrasting continuous and threshold accounts. *Psychonomic Bulletin & Review*, *28*(4), 1112–1130. https://doi.org/10.3758/s13423-020-01862-0
